# SANS (USH1G) regulates pre-mRNA splicing by mediating the intra-nuclear transfer of tri-snRNP complexes

**DOI:** 10.1101/2020.11.11.378448

**Authors:** Adem Yildirim, Sina Mozaffari-Jovin, Ann-Kathrin Wallisch, Jessica Ries, Sebastian Ludwig, Henning Urlaub, Reinhard Lührmann, Uwe Wolfrum

## Abstract

Splicing is catalyzed by the spliceosome, a compositionally dynamic complex assembled stepwise on pre-mRNA. We reveal links between splicing machinery components with the intrinsically disordered ciliopathy protein SANS. Pathogenic mutations in *SANS/USH1G* lead to Usher syndrome – the most common cause of deaf-blindness. SANS functions have been associated with cytoplasmic processes so far. Here, we reveal molecular links between SANS and pre-mRNA splicing catalyzed by the spliceosome, a compositionally dynamic complex assembled stepwise on pre-mRNA in the nucleus. Here, we show SANS associates with Cajal bodies and nuclear speckles, where SANS interacts with components of spliceosomal sub-complexes such as SF3B1 and the large splicing cofactor SON but also with PRPFs and snRNAs related to the tri-snRNP complex. SANS is required for the transfer of tri-snRNPs from Cajal bodies to nuclear speckles. SANS depletion alters the kinetics of spliceosome assembly, leading to accumulation of Complex A. SANS deficiency and *USH1G* pathogenic mutations affects splicing of genes related to cell proliferation and USH. Thus, we provide the first evidence that splicing deregulation may participate in the pathophysiology of Usher syndrome.

## Introduction

Pre-mRNA splicing is a fundamental process in eukaryotic cells that occurs in the nucleus. Splicing results in the removal of introns and the ligation of exons in a nascent precursor messenger RNA (pre-mRNA) to form mature RNA (mRNA) (1). Splicing allows the inclusion/exclusion of exons/introns in the mRNA that can give rise to the synthesis of multiple alternative protein isoforms from a single genomic locus. This process is catalyzed by the spliceosome, a highly dynamic macromolecular complex composed of five small nuclear ribonucleoproteins (U1, U2, U4/U6, U5 snRNPs) and numerous other polypeptides (1, 2). During each splicing cycle, the spliceosome assembles *de novo* on the pre-mRNA by stepwise recruitment and assembly of the pre-formed snRNPs and non-snRNP proteins (3). Process starts with the accumulation of heterogeneous nuclear ribonucleoproteins (hnRNPs) for the splice site selection (complex H), then, U1 bounds to 5′ splice site (complex E) and U2 snRNPs the branch point site of the pre-mRNA, to form the complex A. The U4/U6.U5 tri-snRNP is then recruited to the complex A to form the pre-catalytic spliceosome (complex B). Complex B undergoes conformational and compositional rearrangements, including the release of U1 and U4 snRNPs, yielding the catalytically active complex B*, which catalyzes the first transesterification reaction. After additional rearrangements, complex C is formed catalyzing the second step of splicing which leads to excision of the intron lariat and ligation of the neighboring exons (2,4,5).

The biogenesis of snRNPs is an intricate process that occurs in cytoplasmic and nuclear compartments (6). Newly synthesized U snRNAs in the nucleus are exported to the cytoplasm for additional maturation steps. Subsequently, they are reimported to the nucleus for additional maturation in Cajal bodies, where the final assembly of U2, U4/U6 and U4/U6.U5 snRNP complexes occurs (6–9). Next, mature snRNP complexes are released from Cajal bodies and are transferred to nuclear speckles, where splicing factors are stored until being supplied to active transcription/pre-mRNA splicing sites associated with nuclear speckles (10, 11). Little is known about the molecular transfer between these organelles per se, especially about the transfer of the mature U4/U6.U5 tri-snRNPs from Cajal bodies to nuclear speckles and their subsequent recruitment to pre-spliceosomes.

Conceivably, changes in alternative splicing patterns are important for cell development and differentiation under normal conditions (12, 13). Aberrant splicing patterns and defects in spliceosomal proteins have been implicated in human diseases (13–15). Notably, pathologic mutations in several genes coding for pre-mRNA-processing factors (PRPF) cause *Retinitis pigmentosa* (RP) (16). Here, we demonstrate that defects in a protein related to a syndromic retinal ciliopathy, the human Usher syndrome (USH) affects splicing patterns of genes associated with cell proliferation and USH.

USH is the most frequent cause of inherited combined deaf-blindness, clinically and genetically complex (17, 18); so far, 10 USH genes have been assigned to three clinical types (USH1-3). USH1 is the most severe form characterized by profound hearing impairment, vestibular dysfunction and RP. Myosin VIIa (USH1B) and scaffold proteins such as harmonin (USH1C), whirlin (USH2D) and SANS (USH1G) organize a common USH protein interactome (19). The association of USH proteins with primary cilia, and defects in ciliogenesis and ciliary maintenance caused by their deficiencies characterize the USH disease as a ciliopathy (20–24). Ciliopathies are mainly considered as molecular defects in ciliary modules, such as ciliary transport or signaling (24); however, more recently, ciliopathy genes were also found to be related to the machineries associated with translation and DNA damage-repair processes (25–28). In the present study, we provide the first evidence for molecular and functional links between a ciliary protein, namely the USH1G protein SANS and the pre-mRNA splicing machinery.

The USH1G protein SANS is a scaffold protein harboring several protein-binding domains able to form homodimers (Fig. 1A) (29, 30). Like other USH proteins, SANS is essential for auditory hair-cell development and is part of the signal transduction complex at the tips of stereocilia where it forms high-density protein condensates via liquid-liquid phase separation (31–33). In the eye, the role of SANS and the ophthalmic pathogenesis pathway for USH1G are less clear. SANS is expressed in the retinal photoreceptor cells and glia cells, where it is thought to organize networks of USH proteins which may foster mechanical stabilization of the photosensitive ciliary outer segment (24, 34) and participate in intracellular transport processes (17,21,29,35). Recently, we reported the role of SANS in endocytosis and primary ciliogenesis (21,35,36). Here, we demonstrate the interaction of SANS with key components of the splicing machinery including splicing factor 3B subunit 1 (SF3B1) and SON, a large scaffolding protein known as a splicing cofactor. We show that SANS is involved in the transfer of mature tri-snRNPs from Cajal bodies to nuclear speckles required for the assembly of catalytic spliceosomes. By this mechanism SANS controls splicing of target genes including USH genes. Our results suggest that defective alternative splicing of USH genes may lead to the sensorineural disorders caused by pathogenic variations in the *SANS/USH1G* gene.

**Figure 1.**
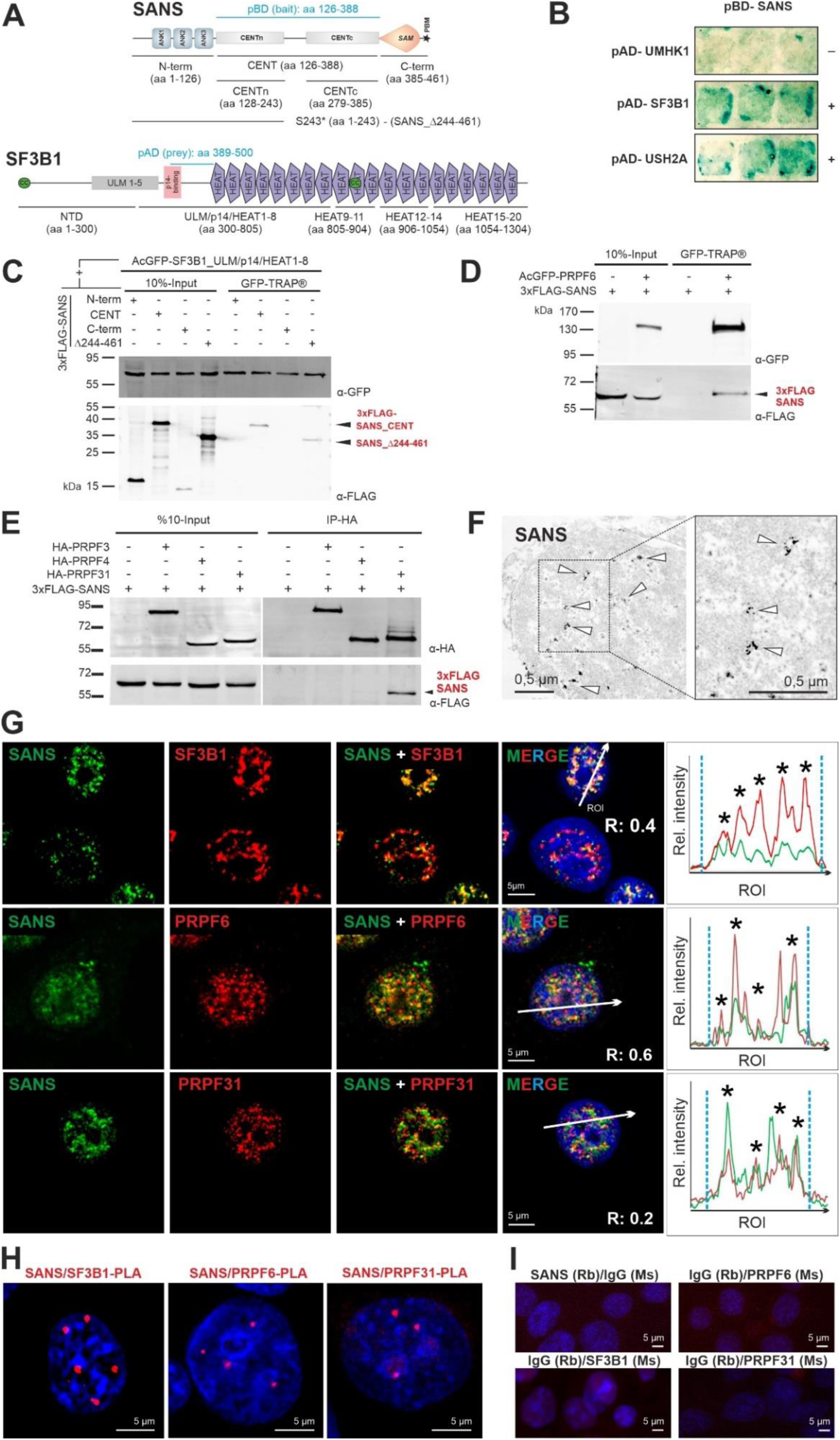
SANS interacts and co-localizes with SF3B1, PRPF6, and PRPF31 in the nucleus. **A)** Domain structures of SANS and SF3B1. SANS is composed of three ankyrin (ANK) repeats, the central domain (CENT), divided into the N-terminal CENTn and the C-terminal CENTc, the sterile alpha motif (SAM) and PDZ binding motif (PBM, *asterisk*). SF3B1 contains the N-terminal domain (NTD) including UHM (U2AF-homology motif) ligand motifs (ULM1-5), p14-binding site, 20 HEAT (**H**untingtin, elongation factor 3/**E**F3, protein phosphatase 2A/PP2**A**, and yeast kinase **T**OR1) repeats and two coiled-coil domains (CC, green). Bait and prey (*light blue*) used for the Y2H screen and deletion constructs are indicated. **B)** 1:1 Y2H ß-galactosidase reporter assay shows binding of pBD-SANS_CENT bait to pAD-SF3B1 prey (aa 389-500) containing parts of the p14 and HEAT1 domain (middle panel) and to pAD-USH2A prey (bottom panel) used as positive control but no binding to U2AF homology motif kinase 1 (UMHK1) (negative control). **C)** Western blot of anti-FLAG IPs from lysates of cells expressing different 3xFLAG-SANS and deletion constructs of acGFP-SF3B1. SF3B1 binds to SANS_CENT and the SANS truncation variant p.S243* (SANS_Δ244-461) reveal CENTn as binding site. **D-E)** Western blot of GFP-TRAP^®^ or anti-HA IPs from HEK293T cells co-expressing 3xFLAG-SANS and GFP- or HA-tagged PRPF31, respectively, demonstrates interaction of SANS with PRPF6 and PRPF31. **F)** Immunoelectron microscopy (anti-SANS RpAb) show theing SANS localization (*arrowheads*) in nuclearplasma of a neuron from the inner nuclear layer of a human retina. **G)** Double immunofluorescence of SANS (MmAb) with SF3B1, and SANS (RpAb), PRPF6, and PRPF31 in nuclei of HEK293T cells counterstained with DAPI. Intensity plots for regions of interest (*white arrows*) indicate co-localization of SANS with SC35 in nuclear speckles (*asterisks*). Blue vertical dashed lines indicate the nucleus extent. **H)** PLAs for SANS in combination with SF3B1, PRPF6 or PRPF31 reveal their interaction in the nucleoplasm of HEK293T cells. **I)** Negative controls for PLAs in H.

## Materials and methods

### Antibodies

As primary antibodies were used: rabbit polyclonal antibodies against SANS (RpAb) (29, 37), anti-SANS (mouse, monoclonal antibody; MmAb) (sc-514418, Santa Cruz), anti-FLAG (F1804, Sigma Aldrich), anti-PRPF6 (sc-48786, Santa Cruz), anti-HA (Roche), anti-GFP (ab6556, Abcam), anti-PRPF31 (PAB7154, Abnova), anti-hSNU114 (38), anti-Coilin (sc-55594, Santa Cruz), anti-SC35 (sc-53518, Santa Cruz), anti-SON (ATLAS, HPA023535), anti-SF3B1 (MmAb) (sc-514655, Santa Cruz), anti-SF3B1 (RmAb)(Will et al 2001), and anti-PRPF38 (homemade). Secondary antibodies were conjugated to the following fluorophores: Alexa488 (donkey-anti-rabbit, A21206, Molecular Probes, Eugene, USA), CF 488 (donkey-anti-guinea pig, 20169-1, Biotrend, Destin, USA), Alexa488 (donkey-anti-mouse, A21202, Molecular Probes), Alexa 555 (donkey-anti-mouse, A31570, Molecular Probes), Alexa 568 (donkey-anti-rabbit, A10043, Invitrogen, Karlsruhe, Germany) CF640 (donkey-anti-goat, 20179-1, Biotrend), Alexa 680 (donkey-anti-goat, A21084, Molecular Probes), Alexa 680 (goat-anti-rat, A21096, Molecular Probes) Alexa 680 (donkey-anti-rabbit, A10043), IRDye 800 (donkey-anti-mouse, 610-732-124, Rockland, Limerick, USA), Abberior STAR Orange (goat-anti-mouse, Abberior, Göttingen, Germany), and Abberior STAR Red (goat-anti-Rabbit, Abberior, Göttingen, Germany). Nuclear DNA was stained with DAPI (4’,6-diamidino-2-phenylindole) (1 mg/ml) (Sigma-Aldrich).

### DNA constructs

3xFLAG-tagged SANS full length and deletion constructs were designed and cloned in house as previously described (29). HA-PRPF31 constructs were kind gifts from Dr. Utz Fischer (University of Wuerzburg, Germany). Deletion constructs of PRPF31 (N-term (aa1-165), C-term (aa 166-499)) were produced by PCR from the HA-PRPF31 plasmid and cloned into the pcDNA3.1 N-HA vector (Thermo Fisher). Deletion constructs of PRPF6 (NTD (aa 1-126), HAT1-6 (aa 293-592), HAT7-13 (aa 593-941)) were cloned into AcGFP-C1 vector (Addgene: 64607). *RON (MST1R)* minigene was a kind gift from Dr. Julian Koenig (IMB-Mainz), and *USH1C* minigene was amplified from the genomic DNA of HEK293T cells and cloned into the pcDNA5/FRT vector (Thermo Fisher). The pcDNA5/FRT-PRPF4 plasmid was constructed by cloning the PRPF4 gene fused to an N-terminal FLAG/HA tag into the pcDNA5/FRT plasmid (Thermo Fisher).

### Yeast two-Hybrid (Y2H)

Yeast two-hybrid (Y2H) screens were performed using the HybriZAP two-hybrid cDNA synthesis kit (Stratagene, La Jolla, USA) as previously described (Maerker et al. 2008). The DNA binding domain (pBD) was fused to the CENT domain of human SANS (amino acids 126-368) (NCBI: NM_173477) and used as bait in a bovine oligo-dT primed retinal cDNA library. The interactions were analyzed by evaluating the activation of the β-galactosidase reporter gene. The interaction between a protein pair was detected by staining the yeast clones in X-ß-gal colorimetric filter lift assay (LacZ reporter gene). For reciprocal 1:1 Y2H assay, bovine SF3B1 (aa389-500) prey fused to the DNA-activation domain (pAD) of the GAL4 transcription factor and human SANS_CENT domain fused to the DNA-binding domain (pBD) of GAL4 transcription factor were used as bait and as prey, respectively. Subsequently, bait and prey were transformed into yeast strains PJ694a and PJ694A, respectively, which were mated and yeast colonies were selected using HIS1, ADE3, LacZ, and MEL1 reporter genes to identify positive interactions confirmed by X-ß-gal colorimetric filter lift assay.

### Immunoprecipitations/pull-down assays

GFP-TRAP^®^ (Chromotek) was performed according to the manufacturer protocol. Briefly, GFP-tagged proteins co-expressed with an indicated 3xFLAG-tagged protein in HEK293T cells. 16 h post-transfection, cells were lysed with Triton X-100 lysis buffer (50 mM Tris–HCl pH 7.5, 150 mM NaCl, and 0.5% Triton X-100) containing protease inhibitor cocktail (PI mix; Roche). Lysates were incubated with the beads for 2h at 4°C followed by several washing steps with 10 mM Tris-HCl pH 7.5, 150 mM NaCl, 0.5 mM EDTA. Bound proteins were eluted with 2x Laemmli buffer, separated by SDS-PAGE followed by Western blotting. FLAG and HA immunoprecipitations (IP) were carried out by using the ANTI-FLAG® (Sigma) and Anti-HA Affinity Gel (Biotool) beads.

### Mass spectrometry analysis

Proteins were separated on 4–12% Bis-Tris-HCl (pH 7.0) NuPAGE polyacrylamide gels (Invitrogen) and stained with G-colloidal Coomassie Brilliant Blue. Entire lanes of the Coomassie-stained gel was cut into 23 slices and processed for mass spectrometry as previously described (39). A comparison of the number of peptides identified for each protein in two samples is a reliable indication of the relative amount of proteins when they are analyzed using identical chromatographic and mass-spectrometric conditions and the same mass spectrometer.

### Cell culture and cell lines

Dulbecco’s modified Eagle’s medium (DMEM) and DMEM-F12 containing 10% heat-inactivated fetal calf serum (FCS) (Thermo Fisher Scientific) were used to culture human HEK293T, HeLa cells, Flp-In-293, and murine IMCD3 cells, respectively. Plasmids were delivered to the cells by using Plus Reagent and Lipofectamine LTX (Invitrogen) following the manufacturer’s protocol. HEK293T cells stably expressing FLAG/HA-tagged human PRPF4 was generated using the pcDNA5/FRT-PRPF4 plasmid and Flp-In-293 cells according to the manufacturer’s protocol (Thermo Fisher). Briefly, Flp-In-293 cells were transfected with a 9:1 ratio of pOG44:pcDNA5/FRT-PRPF4 plasmids using Lipofectamine 2000 (Invitrogen) and 48 h post-transfection, cells were selected in medium containing hygromycin. Stable clones resistant to hygromycin were expanded and tested for stable expression of FLAG-PRPF4 by Western blotting.

### Knockdown and quantitative RT-PCR

siRNAs against human SANS (IDT), PRPF6 (Sigma), PRPF31 (IDT), SRSF1 (Sigma) SF3B1 (Sigma) and SON (Dharmacon) were purchased as indicated in Table S3. Non-targeting control siRNA (siCtrl) was purchased from IDT. For knockdowns, HEK293T and HeLa cells were transfected with a final concentration of 20 nM siRNAs using Lipofectamine RNAiMAX (Invitrogen) following the manufacturer’s protocol. Knockdowns were validated by quantitative PCR (qPCR) using gene specific primers (Table S6) and normalized to *GAPDH*. SANS siRNA knockdown efficiency was also validated by Western blotting and immunofluorescence analysis in HEK293T and HeLa cells (Fig. S3).

### Immunocytochemistry and light microscopy

Cells were fixed with methanol for 10 min at -20°C or with 4% Paraformaldehyde (PFA) for 10 min at room temperature (RT) and washed with PBS. Permeabilization was performed with 0.1% Triton-X100 (Roth) for 5 min at RT, followed by washes with PBS and blocking for 45 min with 0.5% cold-water fish gelatin, 0.1% ovalbumin in PBS. Primary antibodies were incubated overnight at 4°C, followed by washing with PBS and incubation with secondary antibodies for 1 h at RT. After three washes, specimens were mounted with Mowiol 4.88 (Hoechst, Frankfurt, Germany) and imaged using a Leica DM6000B microscope (Leica, Bensheim, Germany). For STED microscopy, we applied secondary antibodies from Abberior and imaged specimens on an Abberior 3D-STED Facility Line microscope (Abberior, Göttingen, Germany).

### Image processing and quantifications

Image processing was performed using a Leica DM6000B microscope (Leica), Leica imaging and ImageJ/Fiji software (40, 41). Deconvolutions were carried out with the Leica program by using 5-7 Z-stacks with one iteration step and BlindDeblur Algorithm settings. Relative accumulations in Cajal bodies and nuclear speckles were quantified by the Cell Profiler program and the mean and standard deviation values were obtained from three independent experiments. The threshold settings were adjusted as follows: Green Channel: Manual threshold: 0.1, Red Channel: Manual threshold 0.1. The primary object size (Cajal bodies) was set in between 1 and 40 diameters. Pearson correlation coefficient was calculated by Coloc2 plugin of the ImageJ/Fiji program.

### Electron microscopy

For immunoelectron microscopy we applied a previously established pre-embedding labeling technique (42, 43). Specimen were analyzed and documented in a Tecnai 12 BioTwin transmission electron microscope (FEI, Eindhoven, The Netherlands) as previously described. Images were processed using Adobe Photoshop CS (Adobe Systems).

### Proximity ligation assay (PLA)

For *in situ* proximity ligation assay Duolink PLA probes anti-rabbit^PLUS^, anti-mouse^MINUS^, and Detection Reagent Red were purchased from Sigma. PLA was performed on HEK293T cells as previously described (29). Briefly, cells fixed in buffered 2% paraformaldehyde were incubated with the primary antibodies overnight at 4°C, followed by incubation with oligonucleotide-labelled secondary antibodies (“PLA probes”) for 4 h at 37°C. After several washing steps hybridizing connector oligonucleotides were added and ligation was performed for 30 min at 37°C, to form a closed circle template. This was followed by rolling circle amplification for 100 min, addition of fluorescent-labelled oligonucleotides and analysis by fluorescence microscopy. For the negative controls, samples probed with only one protein-specific antibody and paired with either the rabbit or mouse IgG specific oligonucleotide-labelled antibody. Cells were mounted in Mowiol 4.88 (Hoechst, Frankfurt, Germany) and analyzed with a Leica DM6000B microscope (Leica, Bensheim, Germany).

### RNA-fluorescence *in situ* hybridization (*RNA-FISH*)

For RNA-fluorescence *in situ* hybridization, we used previously described snRNA probes 5’-end-labeled with Alexa-488 fluorescent dye (Invitrogen) in HEK293T cells (44, 45). Cells were mounted in Mowiol 4.88 (Hoechst, Frankfurt, Germany) and analyzed with a Leica DM6000B microscope (Leica, Bensheim, Germany).

### Cell proliferation assay

The proliferation levels of the cell were tested WST-1 cell proliferation assay (Roche) kit according to the manufacturer’s protocol. In brief, after the seeding 5×10^3^ cells/well in 96 well plates, siRNA-mediated knockdowns were performed 24 h later as described above. 72 h after the knockdown cells were treated with 10 µl cell Proliferation Reagent WST-1 and incubated for 4 h at 37°C and 5% CO_2_. The metabolic activity of the cells was detected with Vario Skan Flash (Thermo Fisher Scientific) at 460 nm wavelength to compare the difference between control and siRNA knockdowns.

### Analysis of alternative splicing *in vitro*

Total RNAs were isolated and reverse transcribed using the Superscript III enzyme (Invitrogen) following manufacturer’s protocol. For the analysis of alternative splicing events of the minigenes and genes, forward and reverse primers against alternatively spliced exons were used to amplify each splice variant with BioTherm Taq polymerase (GeneCraft) (primer list, Table S6). In brief, 48 h after siRNA transfection, minigenes were delivered to the siRNA-depleted cells using Lipofectamine 2000, and 24 h later, cells were harvested. Levels of splice variants were analyzed by TAPE 2200 capillary electrophoresis (Agilent) using high throughput D1000 ScreenTapes (Agilent). Data values were obtained from the 2200 Tape station controller program. Quantifications of the isoform ratios were carried out as previously described (1). Briefly, the median percent spliced in (PSI) indexes were obtained from biological triplicates of siRNA knockdowns. These values summarized as robust Z scores according to the effects of each siRNA treatment with the statistical tests explained in (1). Quantile-quantile plots (Q-Q plots) and Pearsońs correlation coefficient values were calculated in excel by using scaled Z-scores of each individual group (siPRPF6, siPRPF31 and siSRSF1) versus the siSANS group. Graphs were prepared by plotting siSANS groups against the siPRPF6, siPRPF31 and siSRSF1 groups, individually.

### *In vitro* splicing and spliceosome assembly assays

*In vitro* splicing assays were carried out as previously described (46). Briefly, nuclear extracts were prepared from HEK293T cells treated with control and SANS siRNAs. Radioactive [^32^P]- labelled *MINX* pre-mRNA was incubated with 40% (v/v) nuclear extracts in the splicing buffer containing 20 mM HEPES-KOH pH 7.9, 3 mM MgCl2, 65 mM KCl, 2 mM ATP, and 20 mM creatine phosphate at 30°C for the indicated time points. The isolated RNAs either were separated on a denaturing 15% polyacrylamide gel containing 7 M urea and the signals were detected by autoradiography or analyzed by RT-qPCR. The cq values were normalized to controls to obtain relative fold change of snRNA bindings.

For the analysis of the spliceosomal complex formation, the splicing reactions (20 µl) were stopped at the indicated time points by the addition of 2 µl heparin (5 mg/ml) and were analyzed by 2% native-agarose gel electrophoresis followed by autoradiography. SANS depleted nuclear extracts are prepared from siSANS-treated HEK293T cells.

### Purification of the tri-snRNP complex and rescue experiments

Purification of the tri-snRNP complex from HeLa nuclear extract was performed as previously described (47) with minor modifications. Briefly, the nuclear extract in the Roeder C buffer containing 20 mM HEPES pH 7.9, 1.5 mM MgCl2, 450 mM KCl, 0.2 mM EDTA and 25% sucrose, was centrifuged two times ½ h at 20,000 rpm. Next, 3 ml of the cleared supernatant was loaded onto a 20-50% sucrose gradient in 20 mM HEPES pH 7.9, 150 mM KCl, 5 mM MgCl2 and 0.1 mM EDTA in a Surespin rotor and centrifuged at 30,000 rpm for 40 h at 4^°^C. After gradient fractionation, fractions were analyzed by 4–12% NuPAGE (Invitrogen) and the U4, U5 and U6 snRNAs were detected by SYBR gold (Invitrogen) staining. tri-snRNP containing fractionations were pooled and centrifuged overnight in an S58A rotor at 25,000 rpm at 4^°^C. Pelleted tri-snRNPs were dissolved in G250 buffer containing 20 mM HEPES pH 7.9, 250 mM KCl, 5 mM MgCl_2_, 0.1 mM EDTA. To further purify the tri-snRNPs, the sample was loaded onto a 5-20% Sucrose gradient in G150 buffer in a Surespin rotor and centrifuged overnight at 25,000 rpm at 4^°^C. After analyzing fractions on a 4–12% NuPAGE, tri-snRNP containing fractions were detected by SYBR gold staining. Purified tri-snRNP complex was directly used in the chase experiment at the zero time point.

### Glycerol gradient ultracentrifugation

To analyze the levels of snRNPs by gradient fractionation, 200 µg of the siRNA-mediated SNAS depleted HEK293T nuclear extract or the control nuclear extract were diluted with an equal volume of gradient buffer (G150: 20mM HEPES pH 7.9, 150 Mm NaCl, 1.5mM MgCl2 and 0.5mM DTT) and sedimented on linear 4ml 10-30% (v/v) glycerol gradients in the G150 buffer. The ultracentrifugation was performed in a Sorvall TH-660 rotor for 14 h at 29,000 rpm (114,000 × g) and the gradients were separated into 24 fractions. The levels of snRNPs in each fraction were analyzed by Northern blotting using 5′-end radiolabeled DNA probes against U1, U2, U4, U6 and U5 snRNAs, as previously described (48).

## Results

### SANS interacts with splicing factors SF3B1, PRPF6, and PRPF31

We identified the splicing factor 3B subunit 1 (SF3B1) as a putative SANS binding partner in yeast two-hybrid (Y2H) screens of a retinal cDNA library with the central domain of the *USH1G* gene *SANS* (SANS_CENT, aa 126-388) (Fig. 1A). SF3B1, also known as SF3b155 or SAP155, is a component of the U2 snRNP complex of the spliceosome (49, 50). 1:1 Y2H assays confirmed the interaction of SANS_CENT with the central region of SF3B1 (aa 389-500) (Fig. 1B). Reciprocal co-immunoprecipitations (co-IPs) of recombinant GFP- and FLAG-tagged proteins co-expressed in HEK293T cells confirmed the interaction of both proteins and delineated the SF3B1_ULM/p14/HEAT1-8 domain and the SANS_CENT domain as binding sites (Figs. 1C, S1A). Co-IPs in Fig. S1B demonstrated that SANS does not bind to the GFP control.

Notably, defects in several components of the spliceosomal U4/U6.U5 tri-snRNP lead to RP, the ocular phenotype in USH patients (16). This prompted us to examine whether SANS also interacts with other components of the spliceosome. To this end, we focused on the U4/U6- specific protein PRPF31, the U5 protein PRPF6 as well as PRPF3 and PRPF4 which are all components of the U4/U6.U5 tri-snRNP complex and all linked to RP. Western blot analysis of co-IPs from HEK293T cells co-expressing FLAG-SANS and GFP-PRPF6 or HA-PRPF31, PRPF3 and PRPF4 revealed that both PRPF6 and PRPF31 also interact with SANS through its CENTn domain (Figs. 1D-E and S1C-D′). In contrast, PRPF3 and PRPF4 did not (Fig. 1E).

*In silico* structural analysis of SANS using the online tools FoldIndex© (51) and DisEMBL^TM^ (52) predicted the SANS CENTn domain (aa 120-250) as an intrinsically disordered region (Fig. S1E). Such regions are characteristics of intrinsically disordered proteins (IDPs) which can bind to diverse proteins in a context-specific manner and their weak multivalent interactions are the basis for phase separation in cells (53).

### SANS is a nuclear component that interacts with SF3B1 and PRPFs within the nucleus

Next, we addressed where in the cell SANS interacts with the spliceosomal proteins. Immunofluorescence and immunoelectron microscopy demonstrated the localization of SANS in the chromatin-free nucleoplasma of the nucleus (Fig. 1F). Consistently with the nuclear localization of SANS, *in silico* analysis using NLStradamus (54) predicted a nuclear localization sequence (NLS) in the C-terminus of human SANS (aa436-RKKILGAVRRRR-aa447) (Fig. 1A). This was confirmed by NucPred, which rated SANS with a high score (0.76) as a putative NLS-containing protein (55). Immunofluorescence analysis also revealed the co-localization of SANS with SF3B1, PRPF6, and PRPF31 in the nucleoplasm. This was affirmed by calculated Pearson coefficients above zero (R = 0.4, 0.6, and 0.2, respectively, for SF3B1, PRPF6, and PRPF31) and by fluorescence intensity plots (Fig. 1G).

In addition, proximity ligation assays (PLAs) with SANS-SF3B1, SANS-PRPF6 and SANS-PRPF31 pairs revealed PLA signals in the nucleus for all three combinations (Fig. 1H), which were absent in the negative controls (Fig. 1I). This data demonstrated a close spatial proximity of the probed proteins (∼30-40 nm) indicating the presence of SANS in complexes with SF3B1, PRPF6 and PRPF31. Taken together, our results strongly support the molecular interaction of SANS with the splicing factors SF3B1, PRPF6 and PRPF31 in the nucleus.

### SANS binds to spliceosomal snRNP complexes and pre-mRNA

To test whether SANS interacts with spliceosomal snRNP complexes, we performed co-IPs with nuclear extracts from HEK293T cells expressing FLAG-tagged SANS or PRPF4, a core component of the U4/U6.U5 tri-snRNP in absence or in the presence of an *in vitro* synthetized *MINX* pre-mRNA to induce spliceosome formation (Fig. S2). Initial Northern blot analysis of PRPF4 IPs revealed strong hybridization signals for U4, U5, and U6 snRNAs indicating a strong pull-down of the U4/U6.U5 tri-snRNP by PRPF4 in the absence or in the presence of pre-mRNA (Fig. S2A, B). In contrast, SANS did not significantly pull down any of the U4, U5, and U6 snRNAs in the absence of *MINX* pre-mRNA while it pulled down these snRNAs after inducing spliceosome formation by the addition of *MINX* pre-mRNA under splicing conditions (Fig. S2A, B). To validate latter data, we additionally analyzed the snRNAs in pull-downs of SANS and PRPF4 by RT-qPCR (Fig. S2). Consistently, we observed results similar to the initial Northern blots (Fig. S2C). Quantification of three independent experiments revealed that snRNA binding to SANS was drastically increased und splicing conditions. Appling RT-qPCRs we also amplified significant amounts of *MINX* pre-mRNA from the pull-downs of binding to SANS and PRPF4 when compared to the mock-transfected controls indicating the interaction of SANS with target pre-mRNA during the splicing process (Fig. S2D). Taken together, our *in vitro* data suggests that SANS is not stably associated with snRNPs, but interacts with the spliceosomal complexes assembled on *MINX* pre-mRNA under splicing conditions.

### SANS is present in nuclear speckles and its depletion alters the speckles morphology

Since nuclear speckles are important sites for splicing factor storage and modification (11), we tested whether SANS is present in these nuclear bodies. We co-stained SANS and the nuclear speckle marker SC35, also known as SRSF2. Deconvolution microscopy revealed the overlap of the fluorescent signals for SANS and SC35 in nuclei (Fig. 2A), affirmed by merging fluorescence intensity plots and by a positive Pearson correlation coefficient value (R = 0.5) (Fig. 2A-A′). 3D-STED super-resolution microscopy confirm the close association of both molecules at the nuclear speckles (Fig. 2B) as previously found for other speckle-resident proteins by super-resolution microscopy (56).

**Figure 2.**
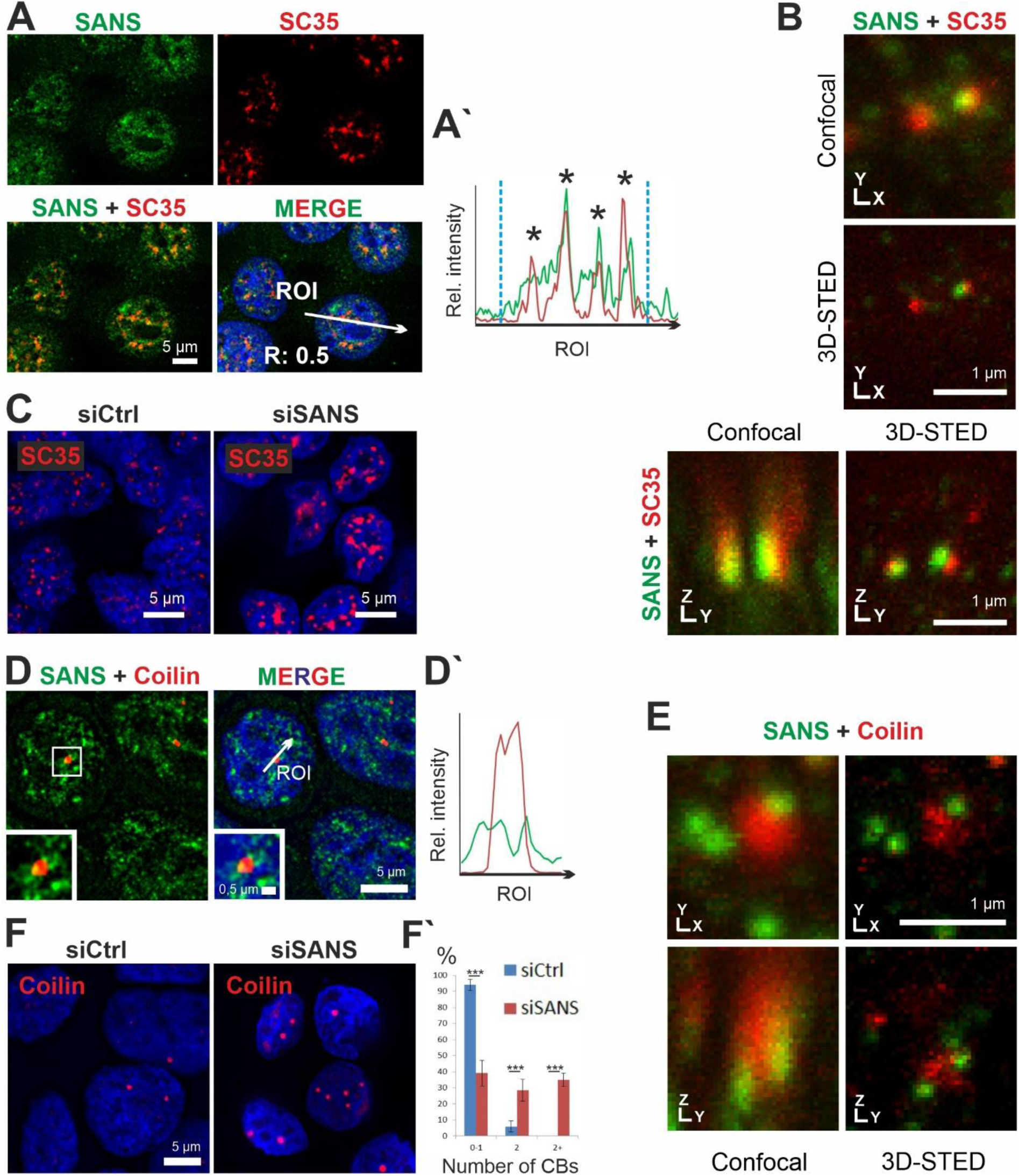
Localization of SANS in nuclear speckles and Cajal bodies. **A)** Immunofluorescence staining of SANS (green) and the nuclear speckle marker/spliceosomal protein SC35 (red) in nuclei of HEK293T cells counterstained with DAPI (blue). **A′)** Intensity plots for the regions of interest in A (*white arrows*) indicate co-localization of SANS and SC35 in nuclear speckles (*asterisks*). Blue vertical dashed lines indicate nuclear extensions. Positive Pearson′s correlation coefficient (R) supports co-localizations of both proteins. **B)** 3D-STED microscopy images of SANS and SC35 co-stained speckles. XY and YZ axes demonstrate close association of SANS with SC35 in nuclear speckles. **C)** Immunofluorescence of SC35 (red) in siRNA-treated HEK293T cells counterstained with DAPI (blue). Nuclear speckles size increases in SANS-depleted cells (siSANS) comparted with control siRNA-treated cells (siCtrl). **D)** Double immunofluorescence of SANS (RpAb, green) and Coilin (red), a molecular marker for Cajal bodies in HEK293T cells counterstained with DAPI (blue). **D′)** Fluorescence intensity plots for region of interest indicated by *arrow* in D reveals SANS localization at the periphery of Cajal bodies in nuclei. **E)** 3D-STED microscopy images of SANS (RpAb, green) and Coilin (red). XY and YZ axes demonstrate localization of SANS at the periphery of Cajal bodies. **F)** Immunofluorescence of Cajal bodies by anti-Coilin (red) in SANS-depleted HEK293T cells. **F′)** Quantification of the number of Cajal bodies (CBs) in SANS-depleted cells; graph shows the percentage of cells with 0-1, 2 or more CBs. SANS depletion increases the number of Cajal bodies in nuclei (In 4 independent experiments 200 and 160 cells were counted in siCtrl and siSANS treated cells, respectively.). ***: p values < 0,001.

The presence of SANS at nuclear speckles prompted us to analyze the effect of SANS depletion on nuclear speckles. To this end, we depleted SANS in HEK293T cells using a validated SANS siRNA (siSANS) and immunostained cells for SC35 (Fig. 2C and S3A-D; Table S1). Confocal microscopy revealed significantly increased and diffused SC35 positive speckles when compared with controls. To exclude off-target effects and cell specificity, we performed the same experiment with an additional siRNA (siSANS #2 recognizing a different region of SANS mRNA) and in HeLa cells as a second cell line. These showed similar diffused speckle formation (Figs. S3D and S4A). This data demonstrated that SANS is present in nuclear speckles where large amounts of the spliceosome components are stored until they are supplied to active transcription/splicing sites or for subsequent post-transcriptional splicing (57, 58).

### SANS is also located at the periphery of Cajal bodies and its depletion increases the abundance of Cajal bodies

PRPF6 and PRPF31 are involved in both the U4/U6.U5 tri-snRNP assembly and its integration into complex A to form pre-catalytic spliceosomes (59). The assembly and maturation of the tri-snRNP complex occur in separate membrane-less sub-nuclear compartments, called Cajal bodies (4). After maturation, tri-snRNP complexes are released from Cajal bodies to join complex A forming complex B at the active splicing sites (4,44,60). To investigate a possible role of SANS for the assembly of the tri-snRNP complex in Cajal bodies, we checked the presence of SANS in Cajal bodies by double immunofluorescence staining of SANS and Cajal body marker Coilin, in HEK293T (Fig. 2D-D′). Image analyses revealed that SANS is localized at the periphery of Cajal bodies. 3D-STED super-resolution microscopy corroborated the localization of SANS at the rim of Cajal bodies (Fig. 2E).

Next, we investigated the role of SANS for the integrity and organization of Cajal bodies by analyzing the effect of SANS′ knockdown on them. Interestingly, siRNA-mediated depletion of SANS led to increased abundance of Cajal bodies both in HEK293T and in HeLa cells (Fig. 2F-F′ and S4B-B′). Consistent with this, knockdown of SANS with a second siRNA (siSANS #2) increased the number of Cajal bodies, despite the milder effect, most probably owing to the lower knockdown efficiency of this siRNA (∼50%) compared with siSANS (Figs. S3A, E-E′ and Table S1).

### Affinity proteomics reveals interactions of SANS with SON in the nucleus

To obtain a comprehensive view of splicing factors associated with SANS in the nucleus, we carried out a proteomic screen by affinity capture of SANS-binding proteins from nuclear extracts. We immunoprecipitated SANS-associated complexes from nuclear extracts of HEK293T cells expressing 3xFLAG-SANS and analyzed the protein content of the immunoprecipitation by mass spectrometry (MS) (Fig. 3A). In the MS data identified 730 proteins from the 3xFLAG-SANS nuclear extract with more than five-fold change in peptide numbers when compared with the mock-transfected control (Table S2A). Gene Ontology enrichment analysis for significantly enriched proteins by DAVID 6.8 (61) revealed “mRNA processing” and “RNA splicing” among the enriched biological processes and “nuclear speckles” as one of the most enriched cellular components (Table S2B-C).

**Figure 3.**
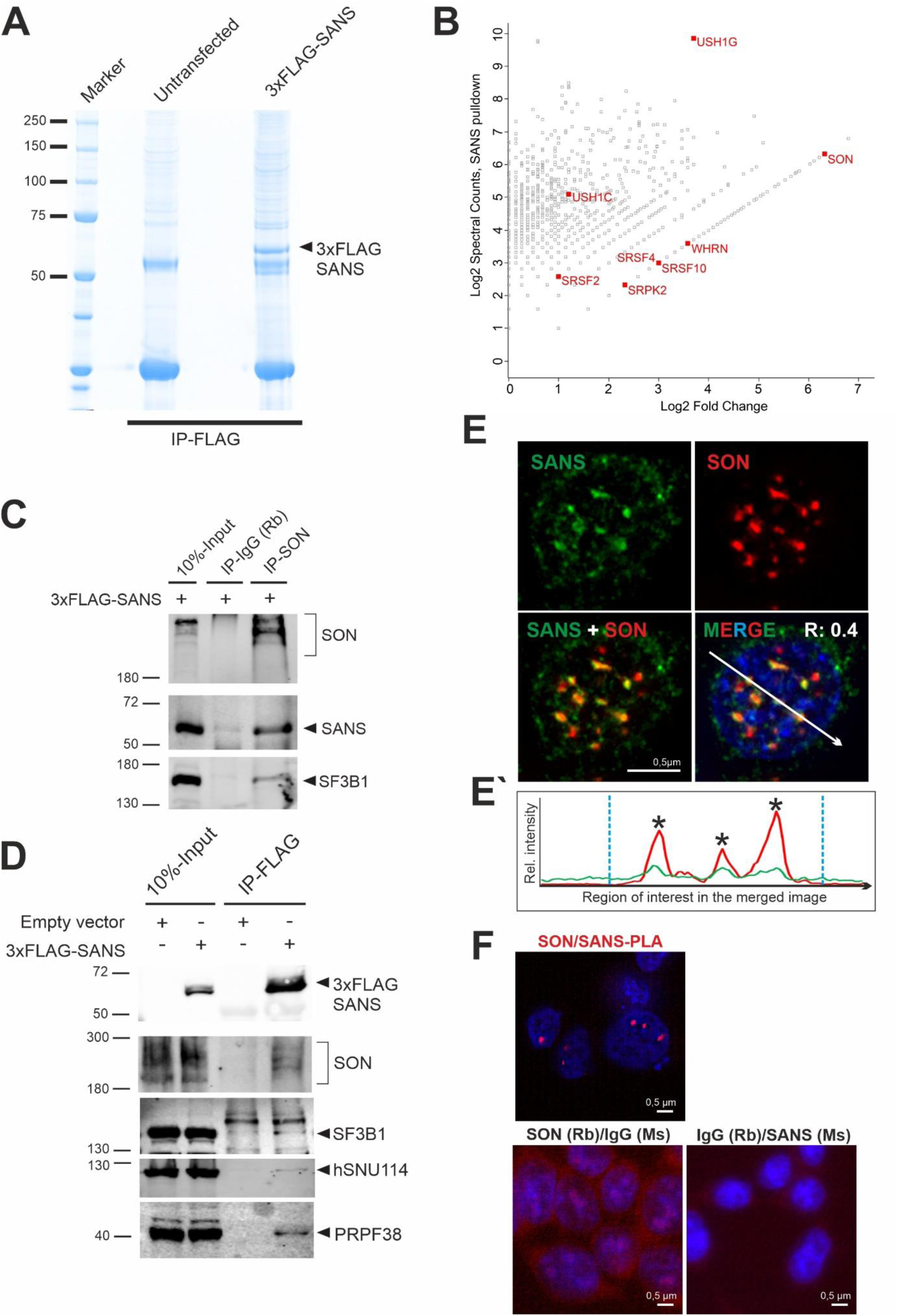
Affinity proteomics and validation of SANS interactions in the nucleus. **A)** Coomassie blue staining of anti-FLAG immunoprecipitated proteins from nuclear extracts of control cells and cells expressing 3xFLAG-SANS, loaded onto a 4-12% NuPAGE. **B)** Proteomic analysis of SANS interaction partners. Graph displays the relative abundance, in logarithmic scale, of proteins identified in 3xFLAG-SANS pull-down from nuclear extract compared with the control. Each dot represents a protein; SR-related splicing factor SON, SR splicing proteins and the SR protein kinase SRPK2 as well as the identified USH proteins are highlighted in red. **C)** Validation of SON-SF3B1-SANS interactions by IP of intrinsic SON in cells expressing 3xFLAG-SANS. SON interacts with SANS and SF3B1. **D)** Western blot analysis of anti-FLAG immunoprecipitations from control and 3xFLAG-SANS nuclear extracts revealed SANS interactions with SON, SF3B1, hSNU114, and PRPF38. **E)** Double immunofluorescence staining of endogenous SANS (MmAb) and SON in HEK293T cells counterstained with DAPI for DNA. Positive Pearson′s correlation coefficient (R) supports co-localization of SANS and SON. **E′)** Fluorescence intensity plots for the region of interest in D (*white arrow*) shows co-localization of SANS with SON in the nucleus. *Blue vertical dashed lines* indicate the extent of the nucleus. **F)** PLAs demonstrate interaction of SANS with SON in cells. Red PLA signals are present in the nucleoplasm in SANS-SON PLAs but absent in the negative controls anti-SANS (MmAb) or rabbit anti-SON with mouse and rabbit IgG antibodies.

Besides known interactors of SANS (e.g. harmonin/USH1C and whirlin/USH2D (Adato et al. 2005; Sorusch et al. 2017)), we found SON – a large Ser/Arg-related splicing factor – as one of the most enriched proteins (Fig. 3B, Table S2A). SON serves as an important scaffold protein for the organization of splicing factors in nuclear speckles and is essential for splicing of genes related to cell proliferation (62, 63). Reciprocal co-IPs from nuclear extract of HEK293T cells expressing FLAG-SANS revealed efficient pull-down of SON with SANS as well as co-precipitation of SANS with anti-SON (Fig. 3C-D). In agreement with the interaction of SANS-SON, immunocytochemistry and PLAs revealed co-localization and spatial proximity of SANS and SON in nuclear speckles *in situ* (Fig. 3E-F). Furthermore, Western blotting of co-IPs for other spliceosomal core proteins demonstrated that SF3B1, hSNU114, and PRPF38 were also co-precipitated with SANS (Fig. 3D). These data corroborate the interaction of SANS to spliceosomal complexes in nuclear speckles, and are in agreement with our results above (Fig. 2C) showing that SANS is essential for the normal morphology of nuclear speckles.

### The tri-snRNP complex accumulates in Cajal bodies of SANS-depleted cells

An increase in the number of Cajal bodies as observed after SANS knockdowns (Fig. 2F) has previously been related to defects in the assembly or in the release of tri-snRNP complexes from Cajal bodies (4, 8). To investigate the role of SANS in the formation of mature tri-snRNPs in Cajal bodies, we first co-stained the U4/U6-specific protein PRPF31 and Coilin in SANS- depleted cells. Confocal microscopy showed the accumulation of PRPF31 in Cajal bodies of SANS-depleted cells when compared with controls (Fig. 4A). Notably, a similar accumulation of PRPF31 in Cajal bodies was previously observed due to tri-snRNP assembly defects after depletion of the U5 protein PRPF6 (4) that we also reproduced (Fig. S4C). In addition, *RNA-FISH* followed by staining for Coilin revealed that U4, U6, and U5 snRNAs accumulated in Cajal bodies in SANS-depleted cells (Figs. 4C-F). Thus, SANS deficiency leads to significant accumulation of tri-snRNP complexes within Cajal bodies.

**Figure 4.**
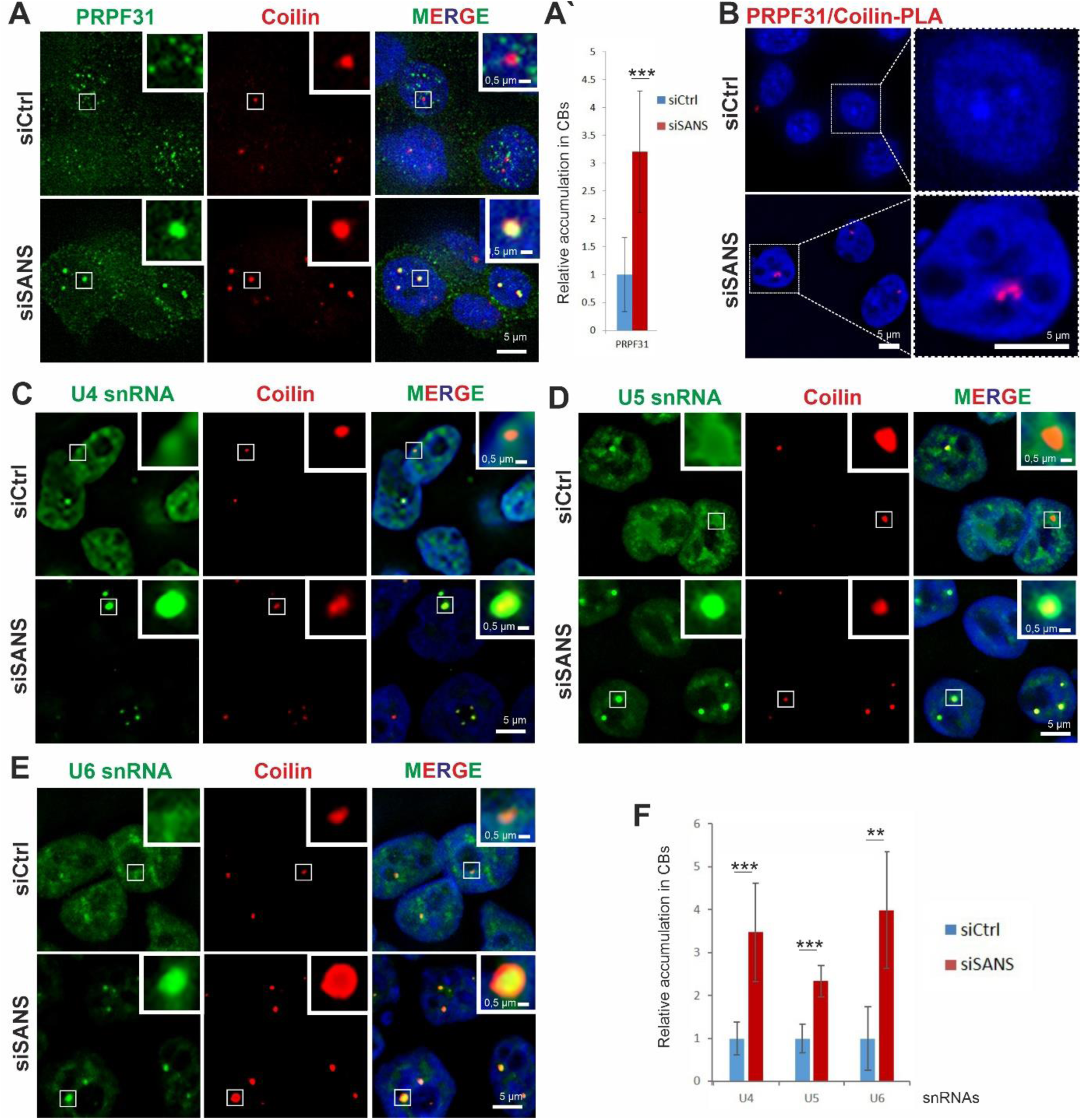
Effects of SANS depletion on Cajal bodies in HEK293T cells. **A)** Double immunofluorescence staining for Coilin (red) and PRPF31 (green) in siCtrl- and siSANS-treated cells. **A**′) Quantification of PRPF31 signal observed in Cajal bodies reveals accumulation of PRPF31 in Cajal bodies (CBs) in SANS-depleted cells. **B′)** PLA shows increased PRPF31-Coilin complexes in CBs of SANS-depleted cells. **C-E)** Analysis of *RNA-FISH* labelling of U4, **C**, U5, **D,** and U6, **E,** snRNAs in CBs (Coilin) in siCtrl- and siSANS-treated cells. **F)** Quantification of accumulation of snRNAs in CBs. SANS depletion leads to the significant accumulation of U4, U5 and U6 snRNAs in CBs. Blue, DAPI counterstaining for nuclear DNA. ***: p values < 0.001, **: p values < 0.01.

Next, we addressed the question whether SANS deficiency also affects the integrity of tri-snRNPs in the nucleus. We depleted SANS in HEK293T cells stably expressing the FLAG-tagged U4/U6 snRNP protein PRPF4 and pulled down the tri-snRNP complex from nuclear extract by immunoprecipitating FLAG-PRPF4. Mass spectrometry analysis (MS) revealed no significant changes neither in the peptide number of components of the tri-snRNP complex nor in its composition when compared with the control (Fig. 5A, Table S3). This was confirmed by glycerol gradient fractionation of nuclear extracts (Fig. S5A). For this, we isolated RNA from the gradient fractions and analyzed their tri-snRNP content by Northern blotting using probes for snRNAs. We detected only a slight increase of tri-snRNPs (fractions 17 and 19, Fig. S5A) in SANS-depleted extracts but we did not observe significant defects in tri-snRNP formation between nuclear extracts of SANS-depleted cells and controls.

**Figure 5:**
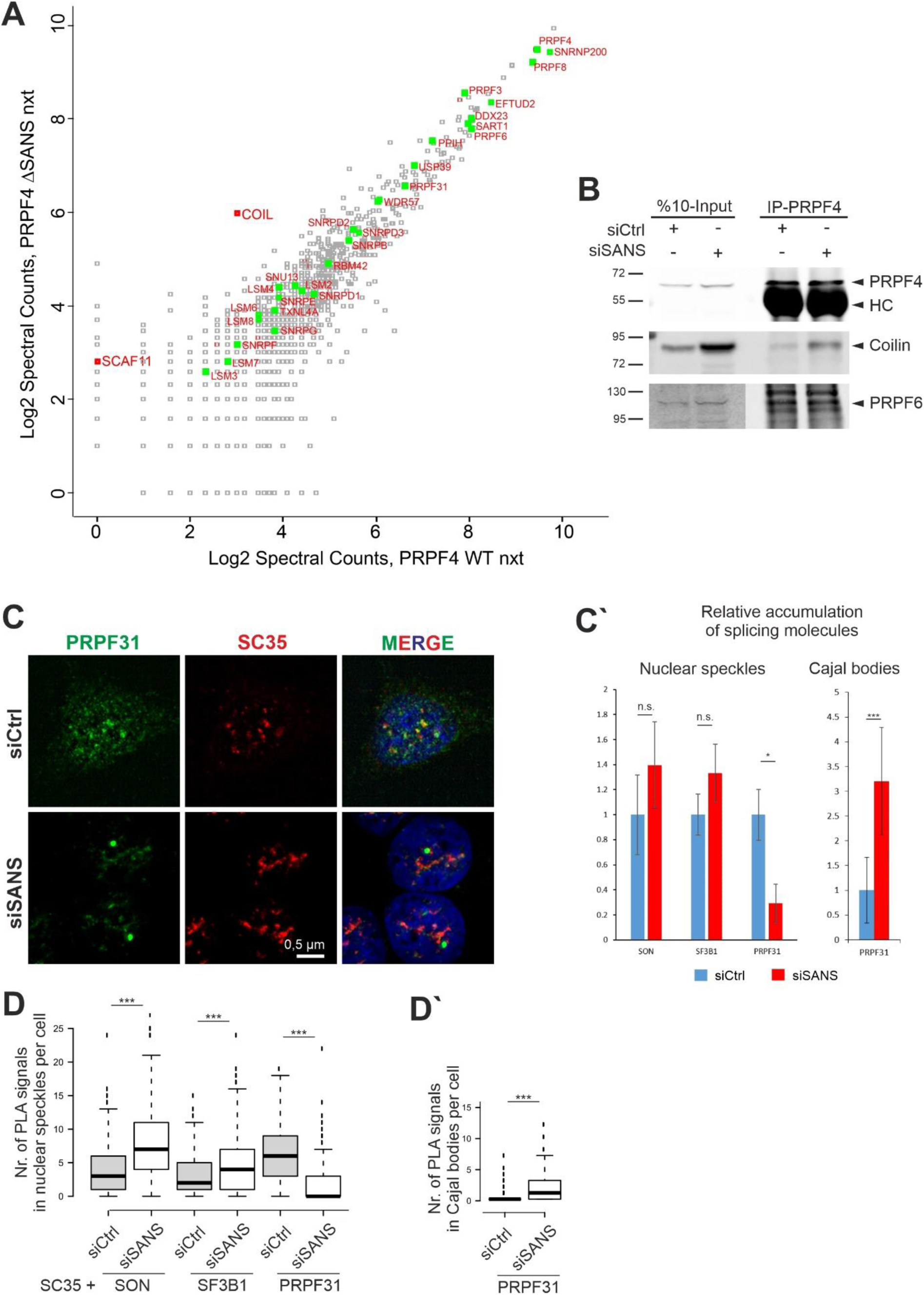
SANS depletion blocks release of tri-snRNP complexes from Cajal bodies and recruitment of mature tri-snRNP complexes to nuclear speckles. **A)** Proteomic analysis of precipitated tri-snRNP complexes via FLAG-PRPF4 from nuclear extracts of control cells and SANS-depleted cells. Graph displays peptide spectral counts for each protein in logarithmic scale. Green: tri-snRNPs; red: Coilin (COIL) and SC35-interacting protein (SCAF11). There was no significant change in the composition of the tri-snRNP in SANS-depleted nuclear extract, but the tri-snRNP was largely bound to the Cajal body scaffold protein Coilin. **B)** Western blot analysis of immunoprecipitation of intrinsic PRPF4 from lysates of HEK293T cells demonstrates stronger interactions of the tri-snRNP protein PRPF4 with Coilin in SANS-depleted cells. SANS depletion does not affect tri-snRNP complex formation visualized by PRPF4 and PRPF6 interaction. **C)** Double immunofluorescence staining reveals reduced PRPF31 staining in SC35 positive nuclear speckles in SANS-depleted cells indicating a defective recruitment of the tri-snRNP complex to nuclear speckles. **C′)** Quantification of immunofluorescence for PRPF31, SON and SF3B1 observed in nuclear speckles and for PRPF31 in Cajal bodies (CB). **D)** Quantifications of PLA signals for SON-, SF3B1- and PRPF31-SC35 show an increase of SON and SF3B1, but a decrease of PRPF31 in nuclear speckles of SANS-depleted cells. **D′)** Quantification of PLA signals for PRPF31-Coilin reveals the accumulation of PRPF31 in Cajal bodies of SANS-depleted cells. : ***: p values < 0.001, *: p values < 0.05.

Interestingly, the Cajal body scaffold protein Coilin was identified by MS as significantly enriched protein in the tri-snRNP fraction precipitated from the siRNA-mediated SANS-depleted nuclear extract (Table S3, Fig. 5A). We validated this result by immunoprecipitations of tri-snRNP complexes using antibodies against Coilin, PRPF4 and PRPF6 (Fig. 5B). While the levels of PRPF4 and PRPF6 were similar between the tri-snRNPs precipitated from control and that of the SANS-depleted cells, we observed an increased binding of Coilin to tri-snRNPs precipitated from the SANS-depleted cells (Fig. 5B).

In conclusion, our data did not provide evidence for a role of SANS in the assembly of the tri-snRNP complexes but indicated that SANS participates in the release of the tri-snRNP complexes from the Cajal bodies.

### SANS depletion blocks the recruitment of tri-snRNPs to nuclear speckles

To investigate the role of SANS in the transfer of tri-snRNPs to nuclear speckles, we first co-stained SC35 with the tri-snRNP protein PRPF31, the U2-specific protein SF3B1 or the splicing cofactor SON in SANS-depleted cells. Notably, the PRPF31 staining in nuclear speckles was significantly decreased in SANS-depleted cells (Fig. 5C) and PRPF31 was mainly localized in Cajal bodies (Fig. 4A). In contrast, SF3B1 and SON staining slightly but not significantly increased in nuclear speckles of SANS-depleted cells (Figs. 5C′ and S5B-C). To examine the formation of spliceosomal complexes in the speckles, we performed *in situ* PLAs to visualize the interaction of SF3B1, SON and PRPF31 with SC35. Interestingly, we found a significant increase in the number of PLA signals for SF3B1-SC35 and SON-SC35 in nuclear speckles of SANS-depleted cells compared with controls (Figs. 5D and S5D). In contrast, PRPF31-SC35 PLA signals were significantly decreased in nuclear speckles of SANS-depleted cells (Figs. 5D and S5D), consistent with our previous immunostaining results: the Cajal body-specific PLA signals for PRPF31-Coilin interaction were significantly increased in these cells (Figs. 4B and 5C′-D′).

Taken together, these results demonstrated that SANS is important for the transfer of the mature tri-snRNP complex to nuclear speckles where tri-snRNPs are stored until being recruited to pre-spliceosomes sites for pre-mRNA splicing. In the absence of SANS mature tri-snRNPs are retained in Cajal bodies leading to the drastic reduction of tri-snRNPs in nuclear speckles.

### SANS depletion reduces kinetics of spliceosome assembly resulting in accumulation of complex A, a delay of complex B formation and inhibition of splicing

Since tri-snRNPs are recruited to the complex A to form fully assembled spliceosomes (complex B), failure in the transfer of tri-snRNP complexes to the nuclear speckles should result in the accumulation complex A, and thus should have an impact on splicing per se. To test this, we examined the effects of SANS knockdown on the spliceosome assembly pathway *in vitro*. We assayed spliceosome complex formation on radiolabeled *MINX* pre-mRNAs using nuclear extracts prepared from control cells (treated with non-targeting control siRNA) and cells depleted for SANS applying siSANS. Splicing reactions were stopped at different time points and spliceosomal complexes were separated by native agarose gel electrophoresis. Figure 6A shows a representative subsequent autoradiography from three independent experiments in total. It indicated substantial changes in the kinetics of the formation of the spliceosomal complexes A and B under SANS-depleted conditions: we observed the accumulation of complex A along with the delayed formation of complex B in SANS-depleted nuclear extracts compared with the control (Fig. 6A-A′, lane 9-14). Quantification of the autoradiograph of the native gel showed that in the control nuclear extract ∼45% of the *MINX* pre-mRNA was present in complex B already after 10 min, and this increased to ∼55% after 30 min (Fig. 6A-A′, lanes 2-4). In contrast, in SANS-depleted nuclear extracts, complex A was accumulated and the formation of complex B was significantly retarded, reaching only ∼35% after 30 min (Fig. 6A-A′, lanes 9-11).

**Figure 6.**
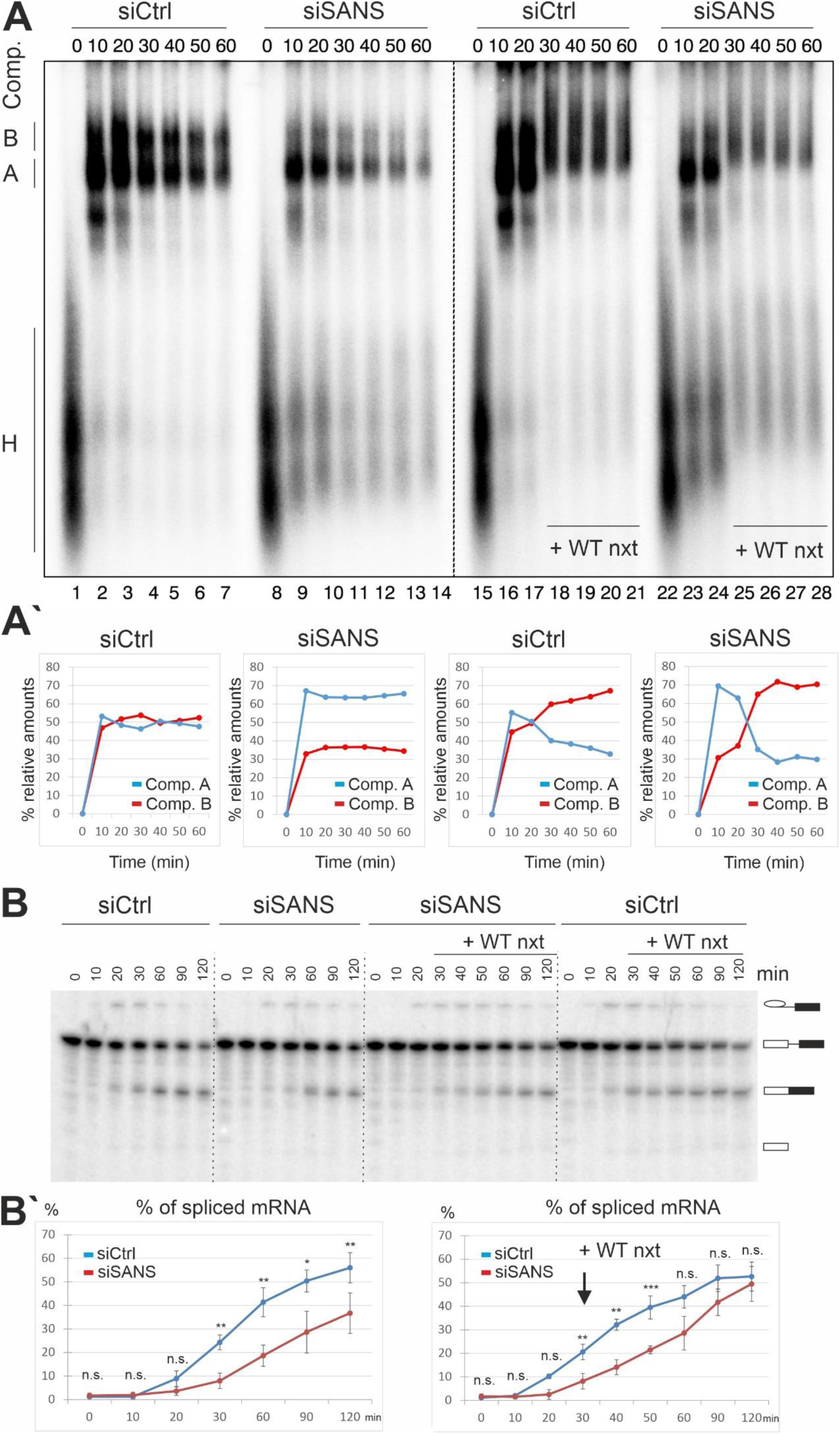
Analysis of the effects of SANS depletion on spliceosome complex formation and splicing. **A)** Autoradiography of the time course of assembly of spliceosome complexes on ^32^P-radiolabelled *MINX* pre-mRNA analyzed by native agarose gel electrophoresis. Splicing reactions were incubated at 30°C for the indicated time points (0-60 min) and stopped by the addition of heparin. Wild type nuclear extract (WT nxt) was added after 20 min to chase the reactions (Lanes: 18-21, 25-28). Positions of each spliceosomal complex are indicated, namely complex H, A and B on the left side of the gel. **A′)** Graphical representation of relative amounts of complex A and B. Each time point in siCtrl and siSANS plots represents an average of three independent experiments with three nuclear extracts separately prepared from cells treated with the indicated siRNAs. The rescue experiments were performed in two independent repeats and were averaged. Depletion of SANS leads to the accumulation of complex A. **B)** *In vitro* splicing of ^32^P radio-labelled *MINX* pre-mRNA with the control and SANS-depleted nuclear extracts. Splicing of pre-mRNA was analyzed by denaturing PAGE followed by autoradiography. **B′)** Quantification of spliced mRNA. Each data point shows an average percentage of spliced mRNA (three independent experiments) for each time point. SANS depletion affects the kinetics of splicing leading to a decreased splicing efficiency (left panel). Addition of WT nxt restores the splicing efficiency (right panel). ***: p values < 0.001, **: p values < 0.01, *: p values < 0.05.

Next, we tested whether the accumulated complex A upon SANS depletion is a functional intermediate or a dead-end complex. For this, we supplemented SANS-depleted nuclear extract with the wild-type nuclear extract (Fig. 6A-A′, lanes 15-28). Autoradiography analysis revealed an effective rescue of the accumulated complex A shifted to complex B (Fig. 6A-A′, lanes 18-21 and 25–28).

Owing to the altered kinetics of spliceosome assembly, we next tested the *in vitro* splicing of pre-mRNA with control and SANS-depleted nuclear extracts. Aliquots were taken at various time points of the *MINX* pre-mRNA splicing reaction followed by RNA isolation and separation of splicing products by denaturing PAGE (Fig. 6B). Quantification of the autoradiographs from three independent experiments revealed that at 120 min only 37% of the pre-mRNA was spliced in siRNA-mediated SANS-depleted extract, while 56% of the pre-mRNA was already spliced in the control reaction at this time point (Fig. 6B′). The impaired splicing was fully rescued by the addition of wild-type nuclear extract to the SANS-depleted extract after 30 min, yielding ∼50% of spliced mRNA (Fig. 6B′). We conclude that the reduced kinetics of spliceosome assembly upon SANS depletion gives rise to impaired pre-mRNA splicing *in vitro*.

### Purified tri-snRNP complex converts the complex A to complex B in SANS-depleted nuclear extract

Next, we tested whether the tri-snRNP complex is the spliceosomal subunit affected in the siRNA-mediated SANS-depleted extract and thus required for converting complex A to the pre-catalytic spliceosome (complex B). To this end, we supplemented SANS-depleted nuclear extracts with purified native tri-snRNPs (Fig. S6). Indeed, the *in vitro* spliceosomal complex formation experiment using radiolabeled *MINX* pre-mRNA demonstrated that while complex A accumulates in SANS-depleted extracts, the addition of the purified tri-snRNPs efficiently rescued the conversion of complex A to complex B. It is worth to mention that the addition of the purified tri-snRNPs did not change the kinetics of the control reaction. Taken together, our results indicated that SANS is important for the supply of the tri-snRNP complexes to the sites of active splicing, required for the assembly of the spliceosome complex B.

### SANS regulates splicing in liaison with pre-mRNA splicing factors in cells

Next, we investigated whether SANS affects splicing of target genes in liaison with core splicing factors in cells. Initially, we analyzed the effects of SANS and PRPF6 depletion on the splicing of *RON (MST1R)* and *USH1C* minigenes expressed in HEK293T cells (Figs. 7A, A′). Analysis of the relative percentage of spliced minigene variants indicated that SANS depletion leads to an accumulation of non-spliced *RON* and *USH1C* pre-mRNAs, by 61% and 29%, respectively, as compared with the control siRNA treated cells. In addition, we observed similar results in PRPF6-depleted cells namely 61% and 20%, respectively.

**Figure 7.**
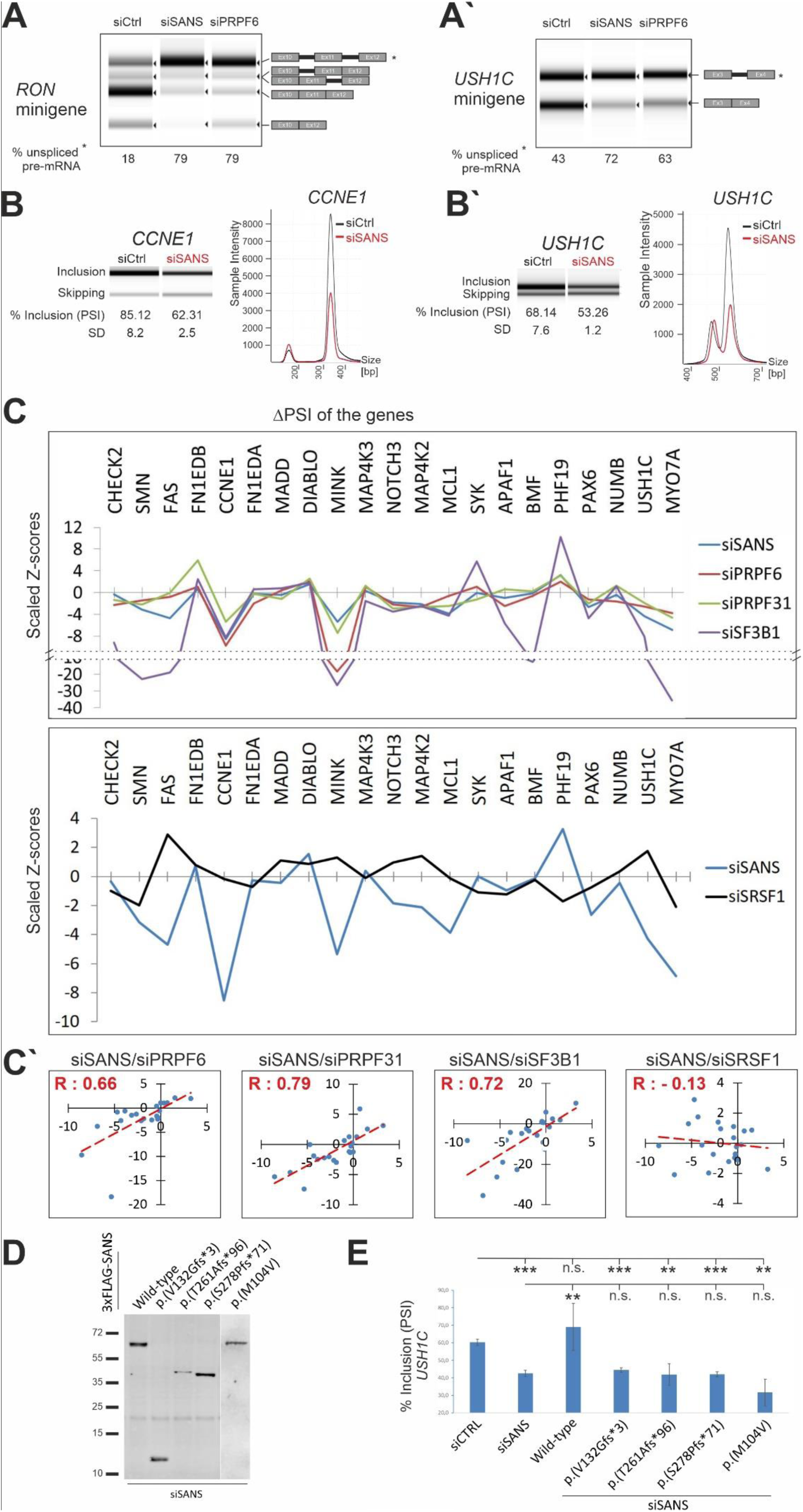
SANS regulates *in vitro* splicing of minigene reporters as well as splicing of genes related to cell proliferation and the human Usher syndrome. **A-A′)** Splicing of *RON* and *USH1C* minigenes in SANS- and PRPF6-depleted cells. Percentage of unspliced mRNA is indicated below lanes. **B-B′)** High throughput capillary electrophoresis analyses of alternatively spliced variants of the *CCNE1* gene (exon 9) and *USH1C* (exon 11) gene after siRNA-mediated depletion of SANS. Relative intensities of the exon included and skipped isoforms obtained from the representative gel images are shown. PSI (% inclusion) and SD indicate, respectively, the percentage spliced-in median and the standard deviation values from three independent experiments. **C)** Graphs showing the splicing perturbation profiles of selected endogenous genes upon knockdown of *PRPF6*, *PRPF31, SF3B1* (upper panel), *SRSF1* (lower panel) or *SANS*. Scaled Z-scores represent changes toward inclusion (>0) or skipping (<0) for each alternative splicing event after siRNA mediated knockdowns. SANS depletion exhibits very similar perturbation profiles to *PRPF6, PRPF31, and SF3B1* knockdowns (upper panel), but to *SRSF1* (lower panel). **C′)** Quantile-quantile (Q-Q) plots are showing positive correlation between SANS, PRPF6, PRPF31 and SF3B1 (Pearson′s correlation coefficient values indicated with positive R values) but not with SRSF1 (negative R value). **D)** Western blot analysis showing the expression of SANS pathogenic variants in the SANS-depleted cells. **E)** Capillary electrophoresis analyses of alternatively spliced variants of *USH1C* (exon 11) gene after siRNA-mediated depletion of SANS following by overexpression of the wild-type and pathogenic USH1G variants. Only wild-type SANS, but none of the pathogenic *USH1G* variants restored the siSANS-induced splicing deficiency in *USH1C*. Relative intensities of the exon included and skipped isoforms obtained from the representative gel images are shown. Quantification of PSI (% inclusion) and SD values shows the percentage spliced-in median and the standard deviation values from three independent experiments. ***: p values < 0.001, *: p values < 0.05.

Next, we examined whether SANS also affects the alternative splicing of intrinsic genes in HEK293T cells. For this, we assessed 19 previously described genes that are alternatively spliced and translated to functional protein isoforms (Fig. 7C) (1). In addition, we included two USH genes *USH1C* and *MYO7A* (USH1B) known to be alternatively spliced (64, 65). After siRNA-mediated knockdowns, we isolated mRNA from cells followed by RT-PCR using splice variant-specific primers to detect alternatively spliced products. As an example, in Figure 7B, we show analyses of the two genes *CCNE1* and *USH1C*, demonstrating a highly significant change in skipping exon 9 or exon 11, respectively (*p* values <0.005).

To depict the splicing perturbation after knockdown of SANS and splicing molecules (PRPF6, PRPF31 and SF3B1 as well as SRSF1) we calculated the percent-spliced-in (PSI) for each gene in three independent experiments (Tables S4 and S5). The scaled Z-scores for the PSIs were plotted in Figure 7C showing the directional change and magnitude of each splicing event. Our results demonstrated that SANS knockdown resulted in a perturbation profile that correlated well with PRPF6, PRPF31 and SF3B1 siRNA-mediated knockdowns (Fig. 7C, upper graph, Fig. S3B). Analyses of Z-scores for SANS, PRPF6, PRPF31 and SF3B1 obtained in three independent experiments revealed positive Pearsońs correlation coefficients between these molecules (R: 0.66, R: 0.79, and R: 0.72, respectively). In addition, quantile-quantile (Q-Q) plots of the Z-scores revealed a positive inclination line (Fig. 7C′) demonstrating similarities between these molecules. As a negative control, we used the knockdown of the serine/arginine-rich splicing factor 1 (SRSF1), which has been previously identified as peripheral component of the spliceosome (1). siSRSF1-mediated depletions gave rise to a different perturbation profile when compared with SANS or knockdowns of the three core spliceosome molecules (Fig. 7C, lower graph, Fig. S3B). Furthermore, no correlation was observed between siSANS and siSRSF1 (negative R: -0.13) (Fig. 7C′). Altogether, our results demonstrated that SANS participates in splicing and that its absence leads to splicing defects similar to essential spliceosome proteins SF3B1, PRPF6, and PRPF31.

### Depletion of SANS and its spliceosomal interaction partners decreases cell proliferation

Our data indicated that SANS is important for alternative splicing of genes related to cell proliferation (Table S5C). Moreover, functional annotation of SANS interaction partners highlighted the cell cycle as one of the enriched pathways (Table S2). We quantified cell proliferation by WST-1 colorimetric assay after siRNA-mediated knockdowns of SANS, PRPF31, SON, and SRSF1, respectively (validation of siRNAs, Fig. S3B). While SRSF1 depletion reduced cell proliferation only to ∼25%, SANS, PRPF31, and SON deficiency, respectively significantly decreased cell proliferation to more than 80% when compared with controls (Fig. S7, Table S5). This data further confirmed that SANS similar to its spliceosomal interaction partners is important for cell proliferation through regulation of alternative splicing of cell proliferation-related genes.

### Pathogenic variants of *SANS/USH1G* affect splicing of the *USH1C* gene

In humans, pathogenic variants of *SANS* lead to USH1G, characterized by combined hearing disorder and RP. Since our previous results showed that SANS deficiency significantly changes the alternative splicing of the *USH1C* gene, we examined whether pathogenic variants of *SANS/USH1G* can also affect the splicing of *USH1C*. We expressed wild-type or four different pathogenic variants of *SANS*, harboring frame-shift or missense mutations, respectively, in cells depleted of endogenous SANS, and analyzed the splicing of *USH1C* (Figs. 7D-E). Our data revealed that the expression of wild-type *SANS* restored *USH1C* splicing defect showing that aberrant splicing caused by SANS depletion can be reversed. In contrast, none of the pathogenic variants of *SANS* was able to restore the splicing defect of *USH1C* in SANS-depleted cells (Figs. 7D-E). This demonstrated that pathogenic variants in *SANS* leading to USH1G disrupt the regulatory function of SANS in splicing of target genes.

## Discussion

We identified unexpected links between the ciliary USH1G protein SANS and the splicing machinery of pre-mRNAs. Splicing is catalyzed by the spliceosome, a highly dynamic macromolecular complex with more than 150-200 proteins contributing to the sequential assembly and regulation of the spliceosome (1, 2). Although previous studies have indicated the participation of SANS in several cellular functions it has not hitherto been related to the splicing machinery. Here, we show that SANS interacts with molecules related to pre-spliceosomal complexes and that it mediates the intra-nuclear transfer of tri-snRNPs between sub-nuclear membrane-less organelles during spliceosome assembly and thereby controls splicing of pre-mRNA (Fig. 8).

**Figure 8.**
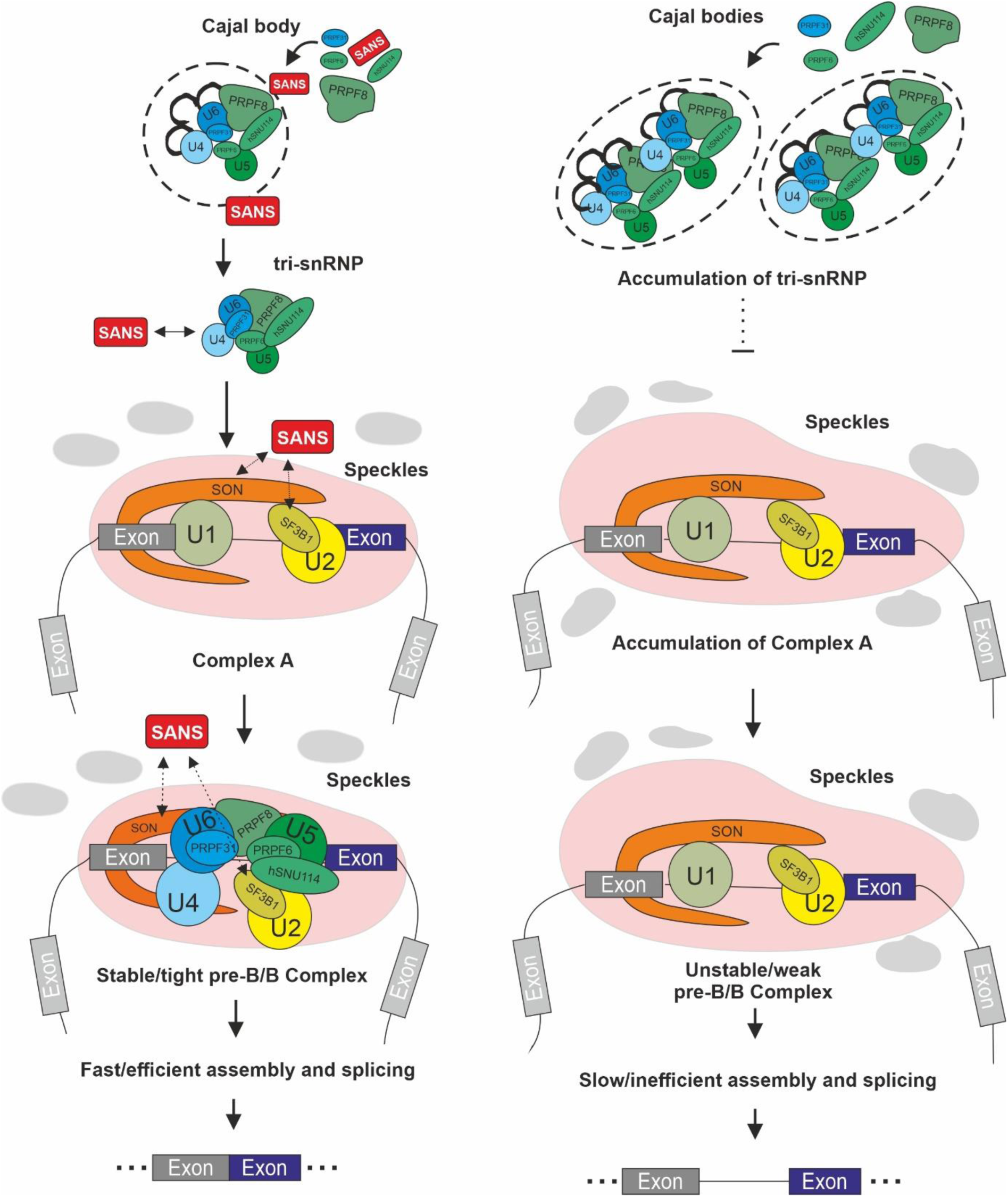
Graphical representation of the role of SANS in the splicing process. In the nucleus the USH1G protein SANS participates at the correct release of tri-snRNP complexes from the Cajal bodies and their transfer to the nuclear speckles for formation of pre-spliceosome B essential and the efficient assembly of active spliceosomes and splicing. In the absence of SANS, assembled tri-snRNPs are not released from Cajal bodies which leads to the accumulation of tri-snRNP complexes in Cajal bodies. tri-snRNP complexes are no longer recruited to the nuclear speckles and the reduced level of mature tri-snRNPs in nuclear speckles retards formation of pre-spliceosome B and B complex, affecting the kinetics and efficiency of pre-mRNA splicing of target genes.

### SANS interacts through an intrinsically disordered region with proteins of the pre-mRNA splicing machinery in membrane-less organelles of the nucleus

SANS is a multivalent scaffold protein and participates in membrane adhesion complexes (29, 35), is associated with microtubule-based intracellular transport (17,30,31,35,66), regulates endocytosis, and is involved in primary ciliogenesis (21). For these functions, SANS interacts with a variety of proteins including several proteins related to the human USH (17,21,29- 32,34,35,66) and other ciliopathies (67). While, PDZ-domain-containing scaffold proteins bind to the type I PDZ-binding motif and the SAM domain at the C-terminus of SANS (21,29,35,68), the majority of interacting proteins bind to SANS via the CENT domain (17,29,30).

In the present study, we demonstrate that core proteins of the pre-mRNA splicing machinery namely SF3B1, PRPF6, and PRPF31 interact with SANS in cells through the N-terminal region of the SANS CENT domain (CENTn). *In silico* structural analysis predicted SANS_CENTn as an intrinsically disordered region (IDR) (Fig. S1E). IDRs can adopt their three-dimensional structures for binding to diverse target proteins in a cell-context-specific manner. This explains the promiscuous interaction of SANS_CENTn with numerous proteins in diverse cellular processes carried out in different cell compartments. Intrinsically disordered proteins (IDPs) are found mainly in macromolecular assemblies and regulate dynamic processes by often transient interactions with target proteins (53, 69). While the binary direct interaction of SANS with SF3B1 was confirmed by both reciprocal co-IPs and Y2H assays, we cannot rule out that the interaction of SANS with PRPF31 and PRPF6 shown by reciprocal co-IPs is indirect via other splicing factors or even RNA. By exploiting these properties, SANS may only transiently or weakly interact with spliceosome molecules, which would explain why it has so far not been found in the proteome of purified spliceosomes.

IDPs are resident proteins of membrane-less organelles such as Cajal bodies and nuclear speckles both of which are the nuclear compartments where SANS is localized and its interaction with the spliceosomal complexes occurs. IDPs are known to drive the formation of both organelles by liquid-liquid phase separation (LLPS) (9,70,71). Recently, it has been suggested that SANS generates protein condensates via LLPS as part of the tip-link complex in stereocilia of inner ear hair cells (33). SANS may change the material properties in the tip-link complex and in the absence of SANS, the tip-link complex of hair cells loose its integrity (32). This feature of SANS is in line with the significant changes in the nuclear speckle morphology which we observed upon SANS depletion (Figs. 2C, S3D and S4A). Therefore, we suppose that the multivalent properties of SANS also promote changes in the material properties of nuclear speckles (72, 73), a feature which remains to be further investigated.

### SANS has no prominent role in the maturation of the tri-snRNP complex in Cajal bodies but participates in the transfer of the assembled tri-snRNPs from Cajal bodies to speckles

Pre-mRNA splicing is a highly dynamic process, characterized by stepwise assembly and release of the spliceosomal U1, U2, U4, U6, and U5 snRNP complexes (2, 74). Northern-blot analysis of SANS pull-downs from nuclear extract did not show enrichment of snRNAs. We concluded that SANS does not directly act as a core splicing factor in the catalytic processes of the splicing. However, under splicing conditions and in the presence of the *MINX* pre-mRNA, SANS could pull down U1, U2, U4, U6, and U5 snRNP complexes suggesting a possible role for SANS in the regulation of splicing activity of at least a subset of genes.

The U4/U6.U5 tri-snRNP complex is a protein-rich particle consisting of more than two dozen polypeptides (2), which assembles and fully matures in Cajal bodies within the nucleus. Failure of tri-snRNP complex maturation, e.g. in the absence PRPF6, leads to the accumulation of incomplete/immature tri-snRNP intermediates in Cajal bodies (for confirmation see Fig. S4C) and the formation of new Cajal bodies (4, 8), which is thought to be a mechanism that compensates for this maturation defect (4, 8). Notably, we observed a similar phenotype in SANS depleted cells: the number of Cajal bodies was drastically increased and SANS deficiency led to the accumulation of the U4/U6-specific protein PRPF31 in Cajal bodies. Furthermore, in SANS-depleted cells, *RNA-FISH* experiments revealed the accumulation of U4, U6, and U5 snRNAs and thus entire tri-snRNP complexes in Cajal bodies. These results may support a role for SANS in the regulation of the tri-snRNP complex assembly which should be evident from the accumulation of the U4/U6 di-snRNP intermediates (44, 46) or from the composition of the isolated tri-snRNP complex. However, analysis of snRNPs by glycerol gradient fractionation as well as the MS analysis in SANS-depleted nuclear extracts showed no significant deficit in the composition of the tri-snRNP complex when compared with those isolated from a wild-type extract (Figs. 5A, B, S5A, S6D and Table S3). In addition, while tri-snRNPs are co-localized with the Cajal body scaffold protein Coilin and thus are located in the core of Cajal bodies, where their assembly is driven (75), microscopy analysis consistently demonstrated the localization of SANS at the periphery of Cajal bodies. Taken together, these findings exclude a key function for SANS in the assembly of the tri-snRNP complex in Cajal bodies. Instead, they suggest a role for SANS in the liberation of assembled tri-snRNPs from Cajal bodies.

### SANS participates in the transfer and recruitment of tri-snRNPs to nuclear speckles fostering the stable assembly of the pre-catalytic spliceosomes

Our microscopy analyses showed that SANS is present in the nuclear speckles – the nuclear compartment where snRNP and non-snRNP splicing factors are stored to be supplied to transcription/splicing sites (11, 58). Our present data demonstrated that the depletion of SANS leads to an increase in size and diffluence of the nuclear speckles, the morphological alterations previously found in splicing-inhibited cells (74). This phenotype was thought to be due to the release of incomplete spliceosomal sub-complexes from the speckles (74) and confirms the role of SANS as a multi-valent intrinsically disordered protein in driving phase separation of nuclear speckles proposed above.

Our *in vitro* interaction studies demonstrated that SANS interacts with several components of the spliceosome indicating that SANS might be involved in the regulation of spliceosome activity. Strikingly our data also revealed the interaction of SANS with SON in nuclear speckles (Fig. 3). SON acts as an important coactivator in splicing and facilitates the formation/stabilization of spliceosomes on weak splice sites by recruiting other SR proteins such as SC35/SRSF2. Through this function, SON contributes to efficient splicing of genes related to cell cycle progression (63, 76). Interestingly, our data showed that SANS also promotes splicing of genes related to cell proliferation and its depletion inhibits cell proliferation. Moreover, we found several SR proteins including SC35/SRSF2 in the SANS affinity proteomics data (Fig. 3B) suggesting that SANS and SON may cooperate as a complex in these activities.

A crucial step before catalytic activation of the spliceosome is the recruitment of the U4/U6.U5 tri-snRNP to complex A, to form the pre-catalytic spliceosomal complex B (60, 77). Our *in vitro* biochemical assays using SANS-depleted nuclear extract and a model pre-mRNA showed the accumulation of complex A corroborating a role for SANS in the formation of stable pre-catalytic spliceosomes (complex B; Fig. 6). Analysis of the recruitment of SON, SF3B1 and PRPF31 to nuclear speckles by double-immunofluorescence staining and PLAs for the interaction with SC35 revealed that SANS is required to recruit the U4/U6.U5 tri-snRNP component PRPF31, but not SON and the U2 snRNP component SF3B1, to nuclear speckles (Figs. 4 and S5). Conversely, SANS deficiency led to slight increases in the staining of SON and SF3B1 in nuclear speckles and to significant increases of PLA signals for SON/SC35 and SF3B1/SC35 complexes, which reflect the accumulation of spliceosomal complexes in nuclear speckles.

Prior to the transfer of the tri-snRNP to nuclear speckles, the assembled complex has to be released from the Cajal bodies. The mechanisms underlying the intra-nuclear transfer processes of snRNPs between nuclear organelles such as Cajal bodies and speckles are yet unknown. Here, we provide an emerging evidence that SANS plays an important role in the release of mature tri-snRNPs from Cajal bodies, and their recruitment to nuclear speckles where assembled spliceosomal complexes are stored for pre-mRNA splicing. Based on our findings described above, we suggest the following mechanism (Fig. 8): the tri-snRNP complexes are assembled in the core of Cajal bodies, diffuse to the periphery where SANS coordinates the release of tri-snRNPs from Cajal bodies for the translocation to nuclear speckles. In the absence of SANS, the release of the tri-snRNP complexes is inhibited and tri-snRNPs accumulate in Cajal bodies (Fig. 4). This leads to deprivation of tri-snRNPs in nuclear speckles required for the formation of the pre-catalytic spliceosomal complex B. Consequently, complex A accumulates leading to significant alterations in splicing kinetics (Figs. 5 and 6). The accumulated complex A in the SANS-depleted nuclear extract is not a dead-end complex. It can be rescued to complex B by the addition of wild-type nuclear extract or isolated tri-snRNP complexes. Furthermore, overexpression of wild-type SANS can rescue the effect of SANS knockdown and increases splicing activity (Fig. 7E).

### SANS is required for correct constitutive and alternative pre-mRNA splicing

The *in vitro* splicing assays with *MINX* pre-mRNA and splicing experiments in cells including minigene reporter splicing assays as well as the analysis of alternative splicing of intrinsic intron-containing genes consistently showed that SANS has a significant impact on constitutive and alternative pre-mRNA splicing. Strikingly, SANS depletion strongly inhibited constitutive and alternative splicing of the *RON* minigene and the constitutive splicing of the *USH1C* minigene (Fig. 7A-A′). In addition, we observed very similar splicing perturbation profiles with positive correlations after the siRNA knockdown of SANS and that of the core components of the spliceosome, namely SF3B1, PRPF6, and PRPF31 (Figs. 7C-C′). In contrast, depletion of SRFS1, a non-snRNP splicing factor known to be involved in splicing regulation (1), gave rise to a different perturbation profile. These findings suggest that SANS interacts not only physically, but also functionally, with core components of the pre-mRNA splicing machinery. Thus, the splicing data obtained in cells are in perfect agreement with the *in vitro* biochemical results, in that both underline the importance of SANS for the delivery of tri-snRNP complexes to nuclear speckles, and thereby for the kinetics of spliceosome assembly.

### *SANS/USH1G* mutations lead to USH due to splicing defects in other USH1 genes

Pathogenic variants in *USH1G* cause human USH1, a ciliopathy characterized by sensory neuronal degenerations leading to profound hearing loss, vestibular dysfunction and vision loss in the form of *Retinitis pigmentosa* (RP) (24). While in the inner ear the essential roles of SANS in hair cell differentiation and the mechano-electrical signal transduction complex can well explain the pathophysiology in USH1G (31–33), the ophthalmic pathogenesis in the retina is still puzzling (78). Links between the USH1G protein SANS – or any other USH1 proteins – and the splicing machinery of cells has not been reported yet. Here, we provide first evidence that perturbation of splicing of target genes can participate in the pathogenesis of the human USH1 phenotype. Our knockdown experiments demonstrate that SANS is involved at essential steps of the splicing process in the nucleus. However, not only depletion of SANS but also USH causing mutations in *SANS/USH1G* lead to splicing defects (Fig. 7E). It is feasible that these mutations alter the physical interaction of expressed SANS variants with components of the splicing machinery, since interaction domains are affected. Alternatively, as pathogenic mutations in *SANS/USH1G* can alter the cellular and nuclear distribution of SANS (29) and thereby SANS might not be any longer recruitable for splicing. In either case, the disruptions in splicing resulting from SANS defects cause splicing errors in other USH1 genes, such as *USH1C* and *MYO7A/USH1B* known to be spliced in the retina (64, 65). It is therefore conceivable that the USH1 phenotype in the retina is not directly caused by SANS, but by mis-splicing of other USH1 molecules. Direct testing of this hypothesis must be deferred for further studies in the future.

### Evidence for a common pathomechanism of SANS and PRPFs deficiencies leading to *Retinitis pigmentosa*

Interestingly, mutations in several pre-mRNA-processing factors of the U4/U6.U5 tri-snRNP – such as *PRPF31* (RP11), *PRPF6* (RP60), *PRPF8* (RP13), *PRPF3* (RP18), *SNRNP200* (RP33) and *PRPF4* (RP70) – also result in retina-specific phenotypes (16,48,79). Our present study shows that deficiencies in PRPFs and SANS cause nearly identical effects on splicing which is in line with significant congruencies between the phenotypes caused by defects in SANS/USH1G and by those in PRPF genes suggesting similar pathomechanisms leading to RP. In this regard, it is noteworthy that both the reduction in expression level due to haploinsufficiency of PRPF31 in RP11 (∼60/∼80% in mRNA, 40-60% reduction in protein level) (Buskin et al, 2018) and the knockdown of SANS/USH1G (∼60/∼80% in mRNA, 40% reduction in protein level) (present study) significantly affected the splicing of target genes. Mono-allelic mutations in the *PRPF31* gene disrupt retina-specific splicing programs and affect ciliogenesis (48, 80). Equally, we have previously shown that SANS depletion alters ciliogenesis (21, 29) and here we demonstrate that SANS depletion affects the splicing programs of target genes, particularly that of *USH1C*, known to be specifically spliced in the retina (64).

Since splicing is a fundamental process that is occurring in many cell types, mutations in tri-snRNP-related genes cause a retinal phenotype only (in the case of RP) or defects in both the retina and the inner ear (in the case of USH). Interestingly, murine animal models for *Prpf* genes and *Sans/Ush1g* do not recapitulate the human ocular phenotype (19,23,81,82). Pre-mRNA splicing factor deficiency may dysregulate specific splicing programs that are known to be specifically implemented in the human retina (83, 84). Recently, it was shown that the truncated pathogenic form of *PRPF31* protein is only expressed in ocular cells derived from RP11 patients, while it was absent in other RP11 patient-derived cells, and that in these non-ocular cells the levels of wild-type protein were nearly equal between normal and patient cells (48). It is thought that the expression of the pathogenic variant protein exacerbates splicing defects, and thus ocular cells are most strongly affected. Nevertheless, it should not be concealed that the PRPF pathogenic variants lead to autosomal dominant RP and USH1G is a recessive disease. However, recent studies show that there are recessive pathogenic variants of splicing molecules such as *CWC27*, *SNRNP200* also causing RP (23, 85). Taken together, there is evidence that the pathomechanisms and pathways leading to RP in patients with *PRPF* or *USH1G* pathogenic variants converge and thereby provide common targets for future therapeutic strategies.

## Conclusion

This study has deciphered the nuclear function of the USH1 protein SANS as a pre-mRNA splicing regulator (Fig. 8). In nucleus SANS acts as a factor required for the correct release from Cajal bodies and recruitment of the tri-snRNP complex to nuclear speckles and thus SANS contributes in the correct assembly of the pre-catalytic spliceosome on target pre-mRNAs. SANS depletion stalls the splicing process at complex A and reduces the kinetics of spliceosome assembly. Consequently, SANS deficiency alters constitutive and alternative splicing of genes related to cell proliferation and to the human USH. Pathogenic variants of *USH1G/SANS* lead to aberrant splice variants of other USH genes which may underlie the pathophysiology of the retinal degenerations associated with USH1G. In addition, our results further support findings of earlier studies, demonstrating that the RP phenotype in the eye can arise from changes in splicing efficiency.

## Supplementary data

### Supplementary figures

Figure S1. Analysis of SANS domains interacting with splicing factors SF3B1, PRPF6 and PRPF31.

Figure S2. Analysis of the interaction of SANS with spliceosomal snRNPs.

Figure S3. Analysis of SANS knockdown on mRNA and protein levels in HEK293T cells.

Figure S4. The effect of SANS and PRPF6 depletion in nuclear bodies in HEK293T and HeLa cells.

Figure S5. SANS depletion affects the nuclear speckle composition.

Figure S6. The purified tri-snRNP complex chases the stalled complex A to complex B in SANS depleted nuclear extracts.

Figure S7. SANS depletion affects cell proliferation.

### Supplementary tables

Table S1. siRNAs and quantifications of knockdowns by qPCR

Table S2. Proteomic analysis of nuclear SANS pull-down and Gene Ontology enrichment analysis

Table S3. Proteomic analysis of FLAG-PRPF4 pull-down from nuclear extract of the PRPF4 stable cell line depleted of SANS

Table S4. TAPE results

Table S5. The effect of SANS knockdown on cell proliferation

Table S6. Primers used in the study

## Author contributions

A.Y. set up and carried out most of the experiments. S.M.J., S.L. and A.Y. set up and carried out the *in vitro* splicing assays and generated stable cell lines. A.K.W. and J.R. helped with several Western blot and immunofluorescence analyses. H.U. and S.M.J carried out the mass-spectrometric analyses. A.Y., S.M.J., and U.W. designed the studies and A.Y. wrote the manuscript under supervision by S.M.J., R.L. and U.W.

## Acknowledgments

We thank to Drs. Utz Fischer, Carlo Rivolta and Julian Koenig for providing several plasmids used in the study; Drs. Cyrill Girard, Dimitry Agofanov and Julian Koenig for valuable discussions; IMB Genomics core facility for the support applying capillary electrophoresis; Mrs. Elisabeth Sehn for her technical expertise with TEM and her valuable help. We also thank to Abberior Company for their valuable support on super-resolution microscopy. This work was supported by GeneRED, IPP Programme Mainz, the FAUN-Stiftung, Nuremberg, and the Foundation Fighting Blindness (FFB; grant no. PPA-0717-0719-RAD).

## Supplementary data

**Figure S1.**
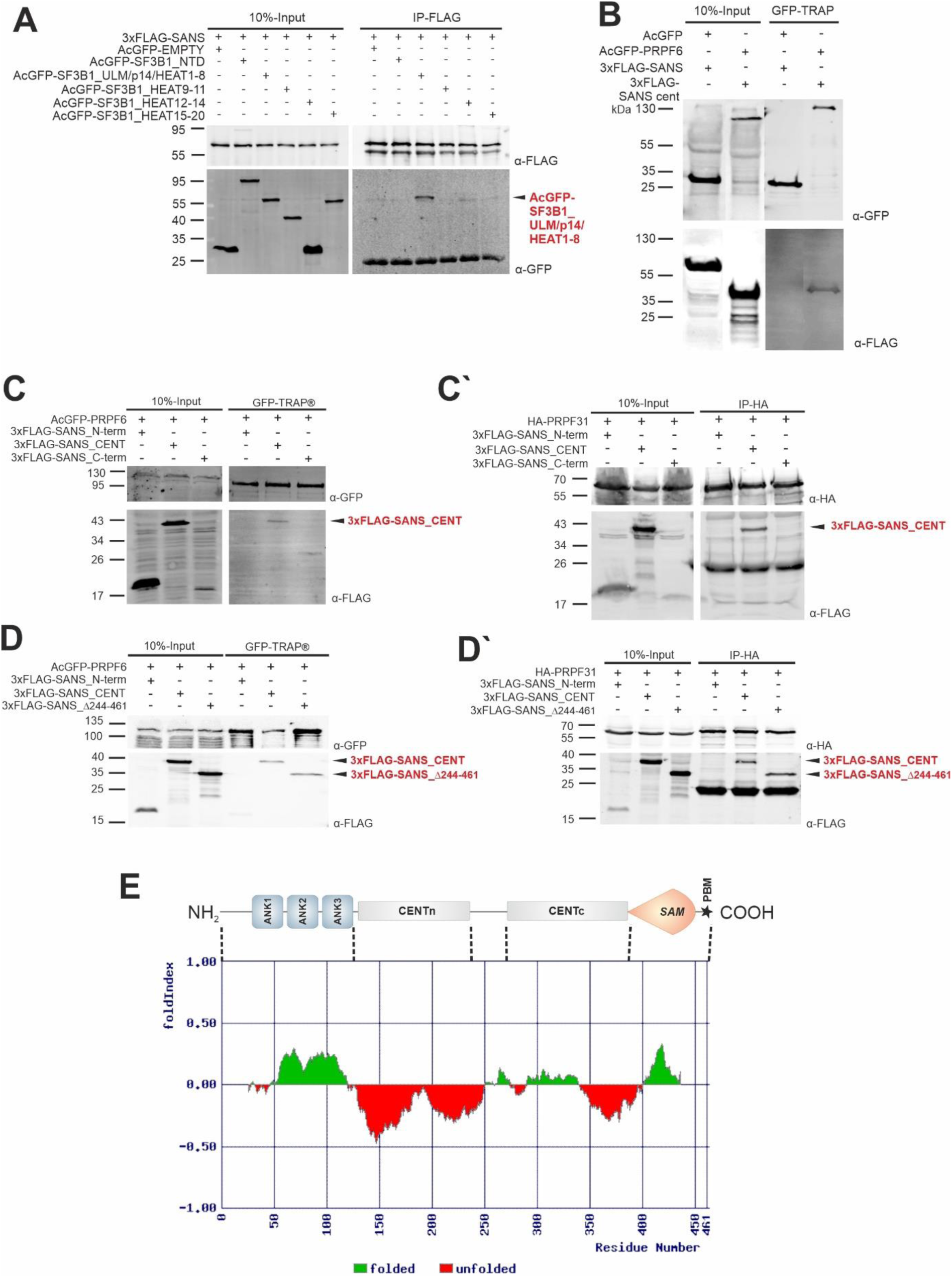
Analysis of SANS domains interacting with splicing factors SF3B1, PRPF6 and PRPF31. **A)** Western blot analysis of anti-FLAG immunoprecipitation from HEK293T cells expressing 3xFLAG-SANS and different truncation variants of SF3B1. SANS interacts with the SF3B1 central region including parts of the ULM, p14-binding and HEAT1-8 domains. **B)** Western blot analyses of GFP-TRAP^®^ pull-downs from cells co-expressing AcGFP-PRPF6 or AcGFP and SANS full length, respectively. Data indicates that only SANS full length interacts with AcGFP-PRPF6 but not with the AcGFP tag or beads of the GFP-TRAP^®^s. **C-C′)** Western blot analyses of GFP-TRAP^®^ or anti-HA pull-downs from cells co-expressing AcGFP-PRPF6 or HA-PRPF31 and SANS truncation variants, respectively. Data indicates that only SANS CENT domain, but not its N-terminal or C-terminal domains, interacts with PRPF6 **(C)** and PRPF31 **(C′)**. **D-D′)** Western blot analyses of GFP-TRAP^®^ and anti-HA pull-downs from cells co-expressing AcGFP-PRPF6 or HA-PRPF31 together with SANS_Δ244-461 lacking the CENTc and downstream domains. Pull-downs show that the SANS CENTn domain is essential for the interactions between SANS and PRPF6 **(D)** and PRPF31 **(D′)**. **E)** Graphical representation of the folding probability for SANS domains computed with the FoldIndex^©^ online prediction tool. CENTn domain is a putative intrinsically disordered region. **A, C-D′:** numbers indicate molecular weights in kDa.

**Figure S2.**
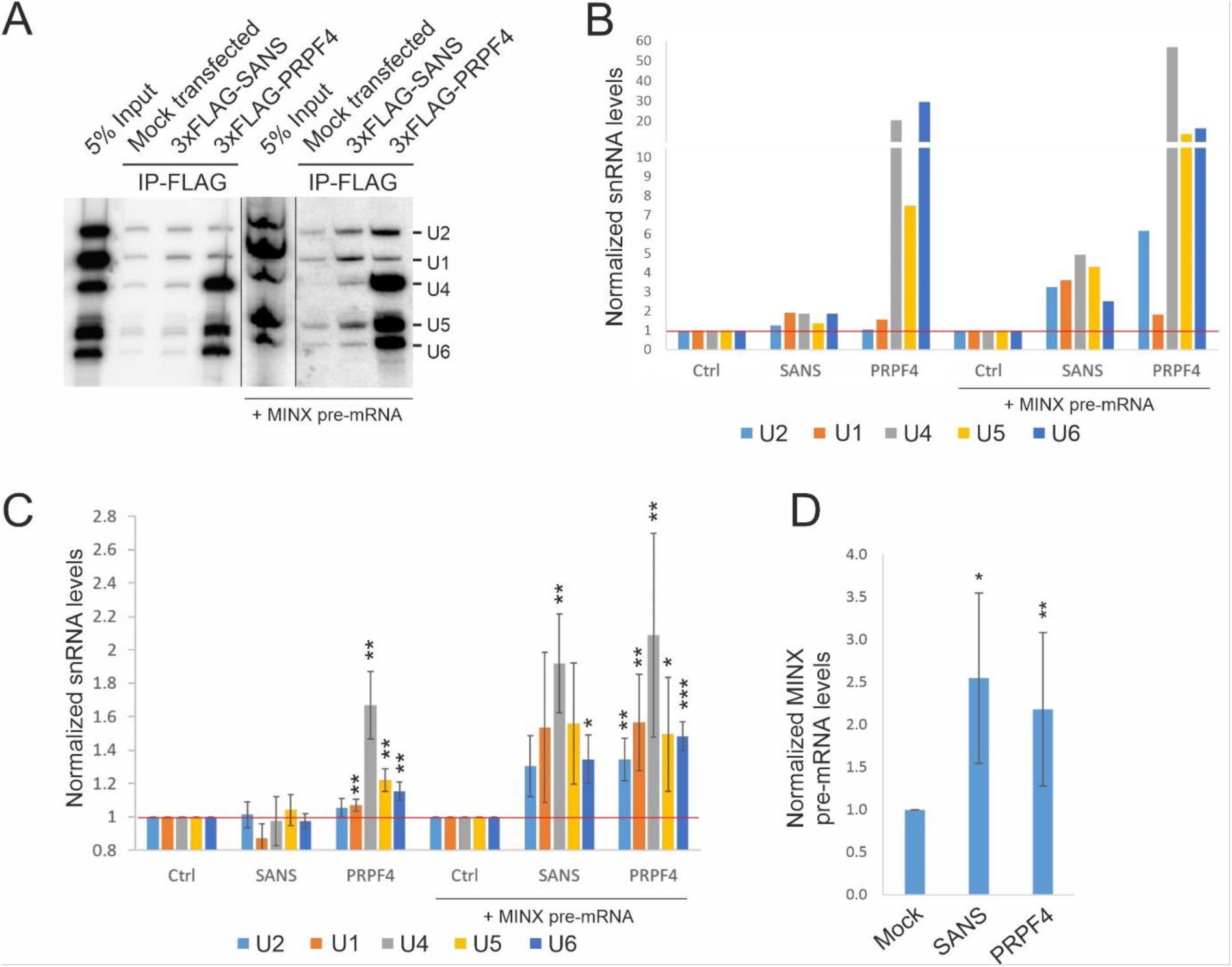
SANS binds spliceosomal snRNPs. **A)** Representative Northern blot analysis of two independent experiments of isolated RNAs from anti-FLAG immunoprecipitations in nuclear extracts of HEK293T cells expressing 3xFLAG-tagged SANS or PRPF4, in the absence or in the presence of *MINX* pre-mRNA (under *in vitro* splicing conditions) applying snRNA probes (U1, U2, U4, U5, and U6). **B)** Densitometry analysis of the Northern Blot in A, indicates that unlike PRPF4, SANS does not bind to U4/U6.U5 tri-snRNPs, but shows binding to all snRNPs under in vitro splicing conditions in the presence of *MINX* pre-mRNA. **C**) Analysis of three independent RT-qPCRs of isolated snRNAs show significant binding of SANS to all snRNPs to under splicing condition in the presence of *MINX* pre-mRNA (N=3) **D)** RT-qPCR analysis showing the binding of SANS and PRPF4 to *MINX* pre-mRNA (N=3). P values: ***: < 0.001, **: < 0.01, *: < 0.05.

**Figure S3.**
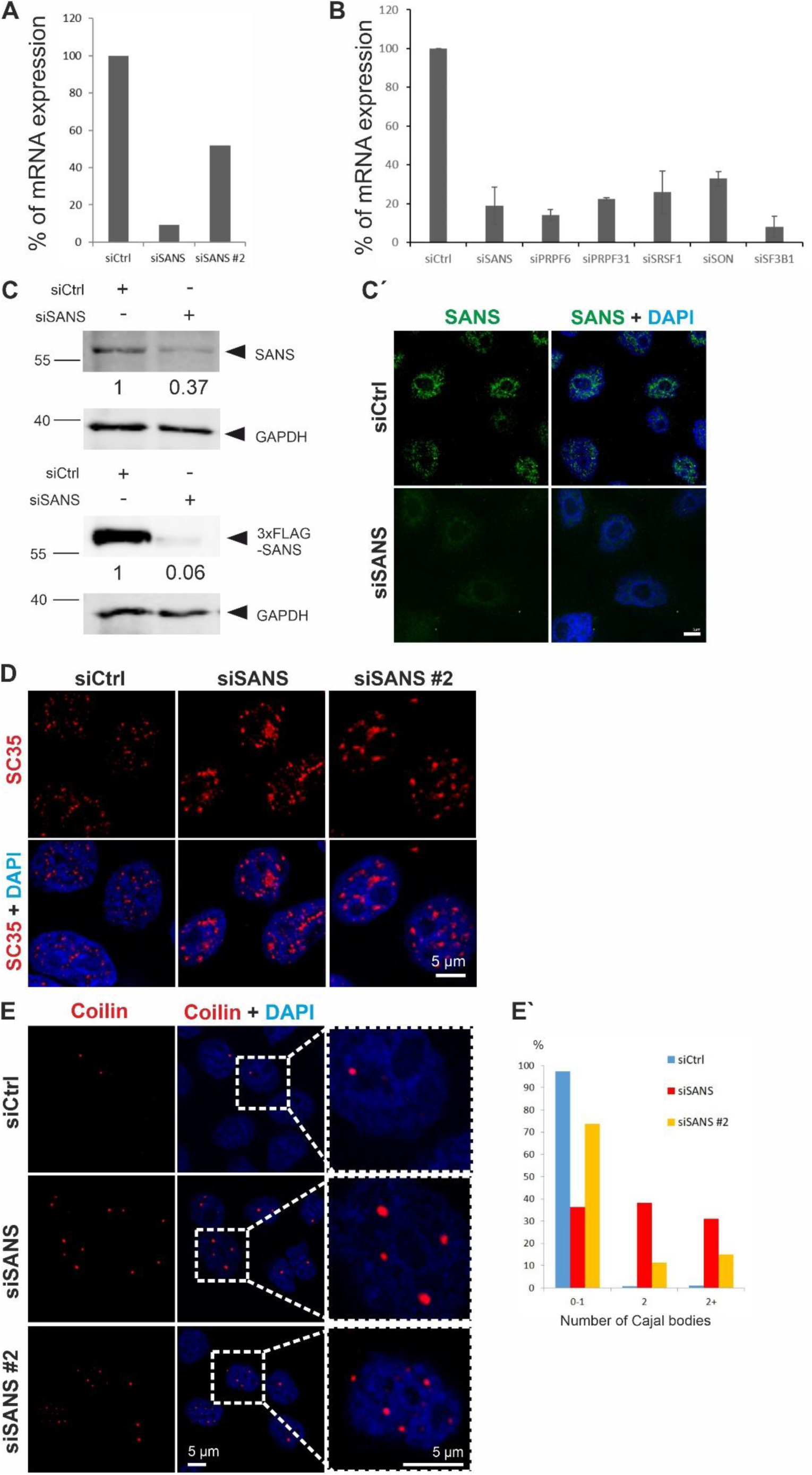
Analysis of SANS knockdown on mRNA and protein levels in HEK293T cells. **A)** Quantitative PCR (qPCR) analysis of SANS mRNA expression in HEK293T cells treated with two different siRNAs (siSANS, siSANS #2). While siSANS leads to ∼90% depletion, siSANS #2 only decreased the level of SANS mRNA to 50%. Thus, we used siSANS for further experiments. **B)** Representative qPCR analysis of mRNA expressions from three independent experiments. The siRNA-mediated knockdowns of SANS, PRPF6, PRPF31, SRSF1, SON, and SF3B1 resulted in ∼80% decrease in mRNA levels. **C)** Western blot analysis of SANS-depleted HEK293T cells. siRNA knockdown of SANS leads to 60% decrease in the protein level of intrinsic SANS (RpAb). In addition, FLAG-SANS cannot be overexpressed in siSANS-treated cells indicating the specificity of the siRNA. **C′)** Immunofluorescence analysis of SANS-depleted HeLa cells. SANS (RpAb) signal disappeared from siSANS-depleted cells. **D)** Immunofluorescence staining of nuclear speckles of SC35 in the cells depleted of SANS applying siSANS or siSANS #2 siRNAs, respectively. Both of siRNA depletions show enlarged nuclear speckle signals. Blue, DAPI counterstaining of nuclear DNA. **E)** Immunofluorescence staining for Cajal bodies (Coilin, red) in SANS-depleted HEK293T cells using siSANS or siSANS #2 siRNAs. **E′)** Quantification of the number of Cajal bodies in SANS-depleted cells; the graph shows the percentage of cells containing 0-1, 2 or more Cajal bodies. siSANS increased the number of Cajal bodies more efficiently than siSANS #2.

**Figure S4.**
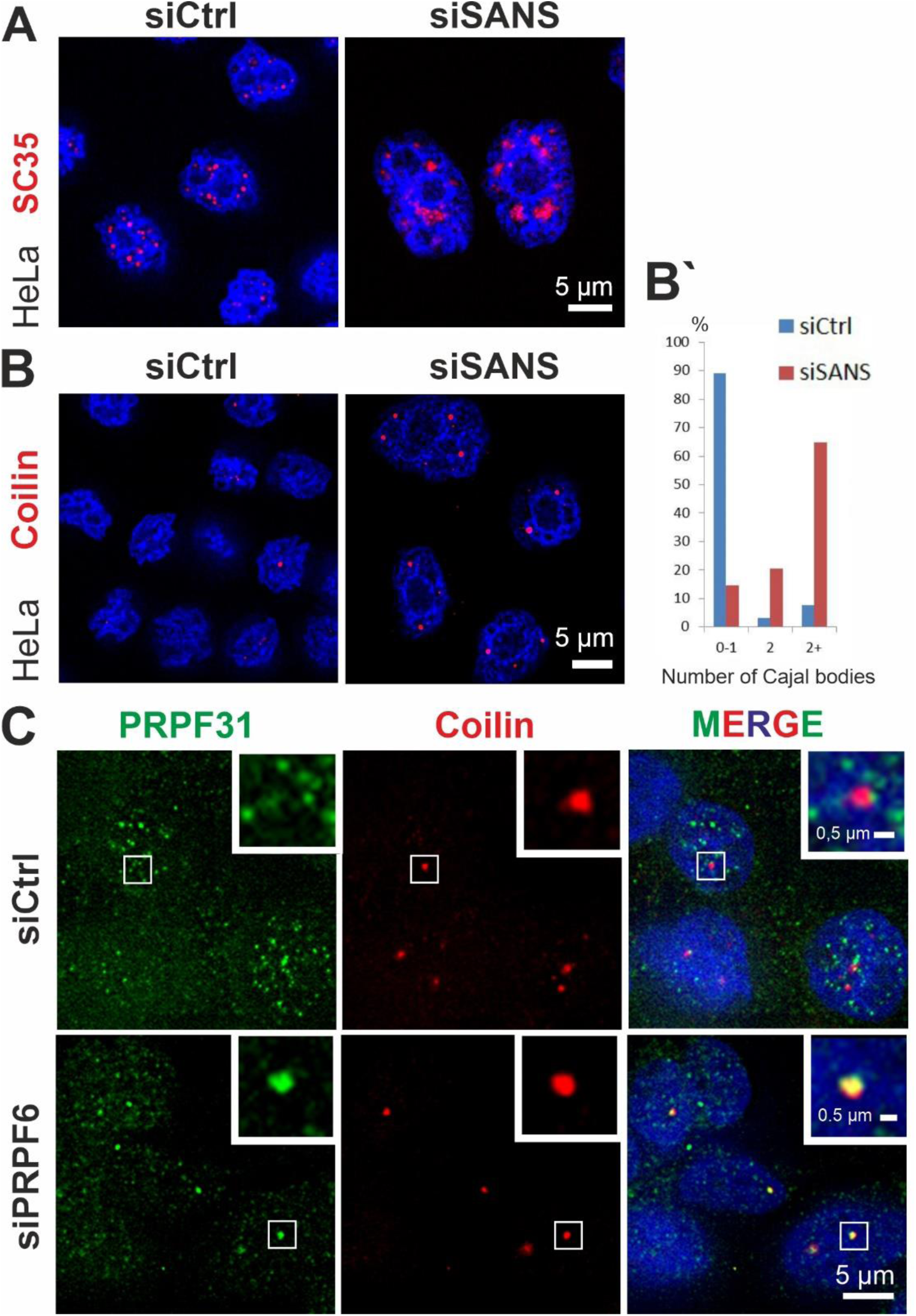
The effect of SANS and PRPF6 depletion on nuclear bodies in HEK293T and HeLa cells. **A)** Immunofluorescence of SC35 (red) in siRNA-treated HeLa cells counterstained with DAPI (blue). Nuclear speckles size increases in cells depleted of SANS (siSANS) compared with control siRNA-treated cells (siCtrl). **B)** Immunofluorescence staining for Cajal bodies (Coilin, red) in SANS- depleted HeLa cells. **B′)** Quantification of the number of Cajal bodies in SANS-depleted cells; the graph shows the percentage of cells containing 0-1, 2 or more Cajal bodies. SANS depletion increases the number of Cajal bodies in the nucleus. **C)** Double immunofluorescence staining of PRPF31 and Coilin in siCtrl- or siPRPF6-treated cells.

**Figure S5:**
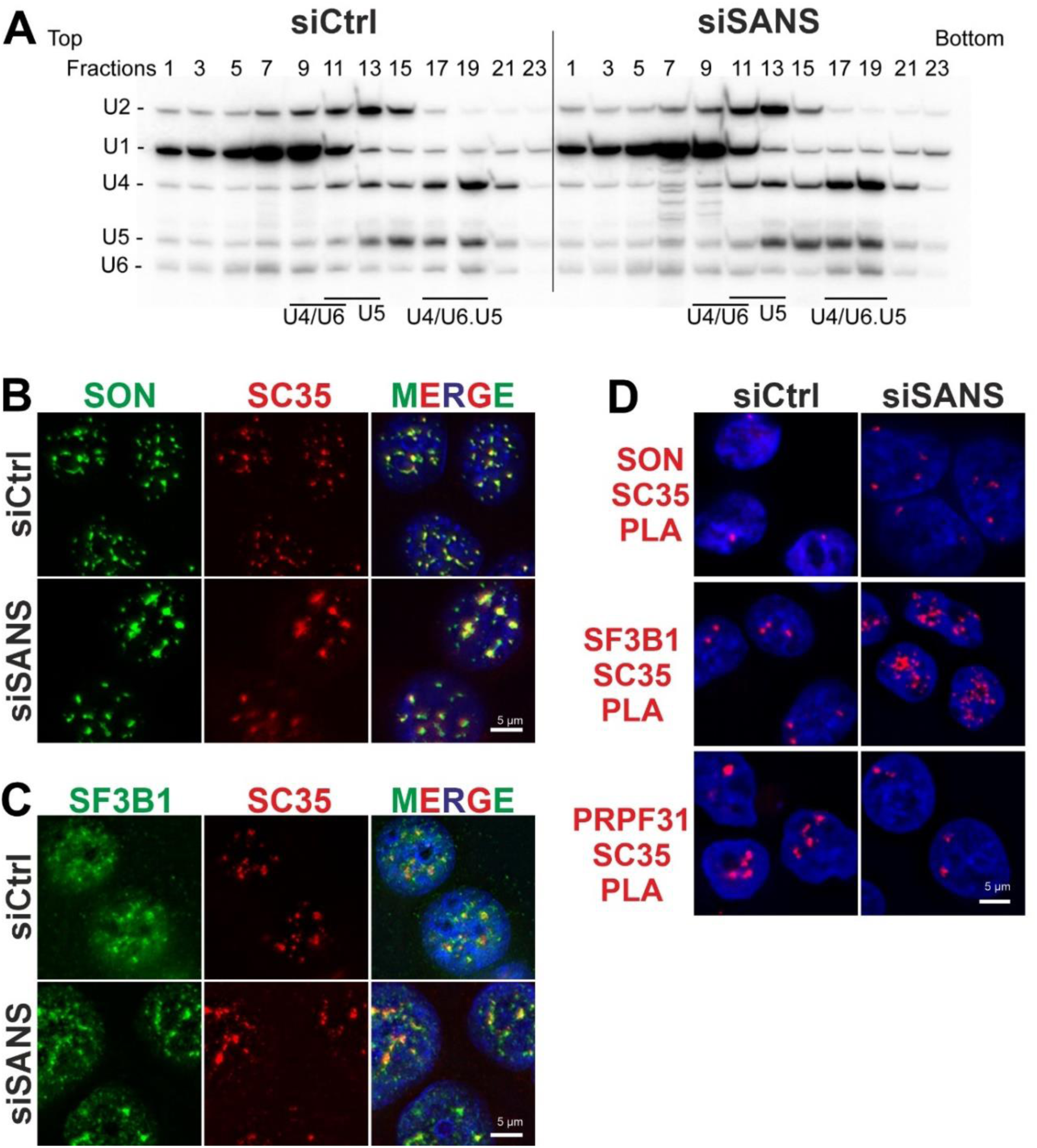
SANS depletion affects the nuclear speckle composition. **A)** Northern blot analysis of snRNAs isolated from a glycerol gradient (10-30%) fractionations of nuclear extracts from siCtrl- and siSANS-treated cells by applying oligonucleotide probes against U1, U2, U4, U5 and U6 snRNAs. The comparison of the distribution patterns of snRNPs across the gradients shows no accumulation of U4/U6 or U5 snRNPs indicating no significant deficit in the formation of the U4/U6.U5 tri-snRNP complex in SANS-depleted nuclear extract (Lane 17-19). However, a slight increase in the amount of tri-snRNPs is detected in the SANS-depleted extract (fraction 17). **B-C)** Immunofluorescence analysis shows increased staining of SON and SF3B1 in nuclear speckles (stained by anti-SC35) in SANS-depleted cells, when compared with siCtrl-treated cells. **D)** PLAs of SON-, SF3B1- and PRPF31-SC35 reveal increase of PLA signals for SON- and SF3B1-SC35 PLA pairs, but a decrease for PRPF31-SC35 confirming accumulations of SON and SF3B1 and the lack of PRPF31 in nuclear speckles of SANS-depleted cells.

**Figure S6.**
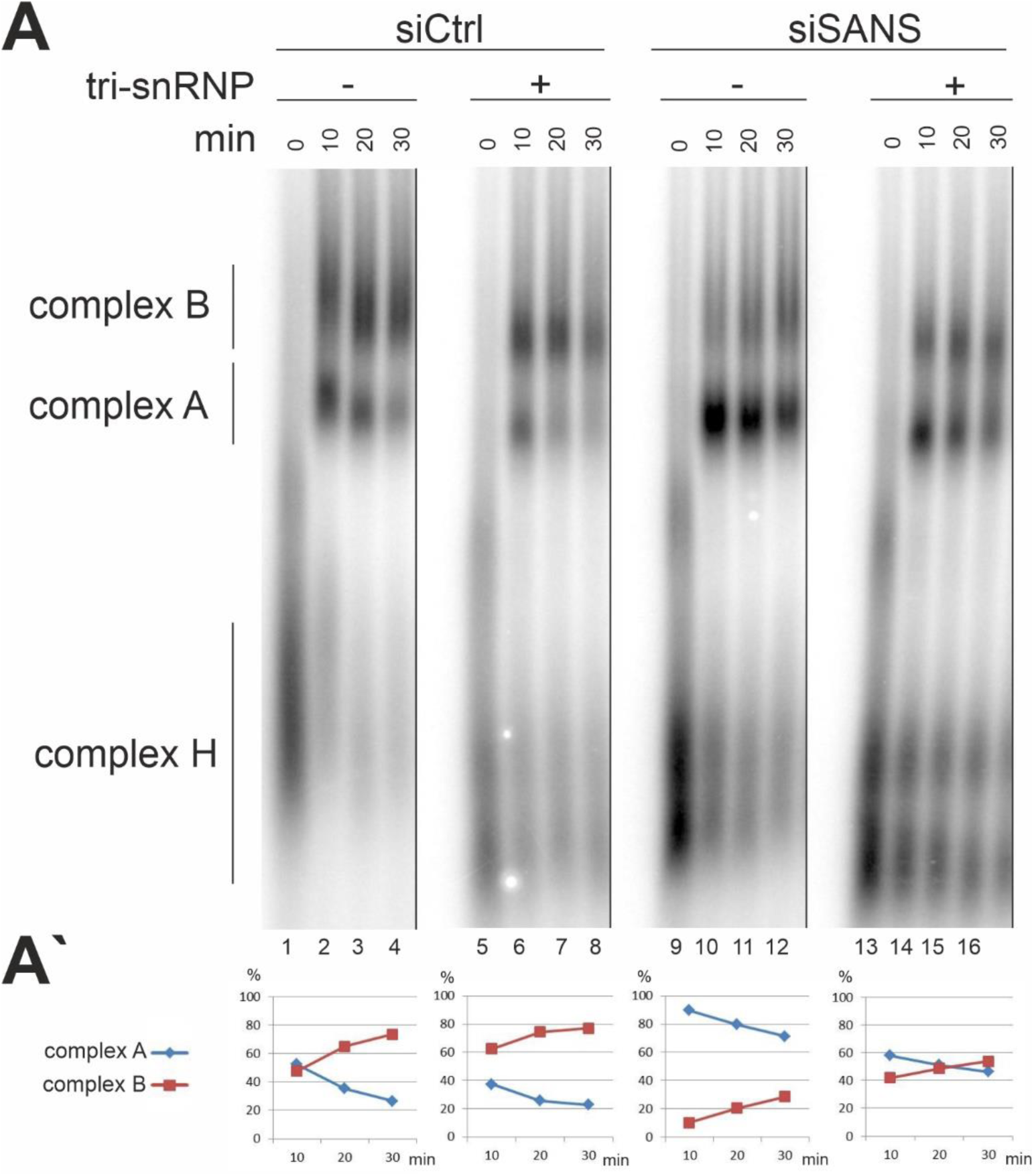
The purified tri-snRNP complex recues the stalled complex A to complex B in SANS-depleted nuclear extracts. **A)** Autoradiography of spliceosomal complexes assembled on ^32^P-radiolabelled *MINX* pre-mRNA in control and SANS-depleted nuclear extracts analyzed by native agarose gel electrophoresis. Splicing reactions were incubated at 30°C for the indicated time points (0-30 min) and stopped by the addition of heparin. Purified tri-snRNP complex is added to the reaction before addition of *MINX* pre-mRNA at 0 min time point (lanes 5-8 and 13-16). Positions of spliceosomal complexes H, A, and B are indicated. **A′)** Quantification of the percentage of spliceosomal complexes A (blue) and B (red) at each time point. In SANS-depleted nuclear extracts addition of purified tri-snRNPs rescues the conversion of the accumulated complex A to complex B (Lane 9-16); N=1.

**Figure S7.**
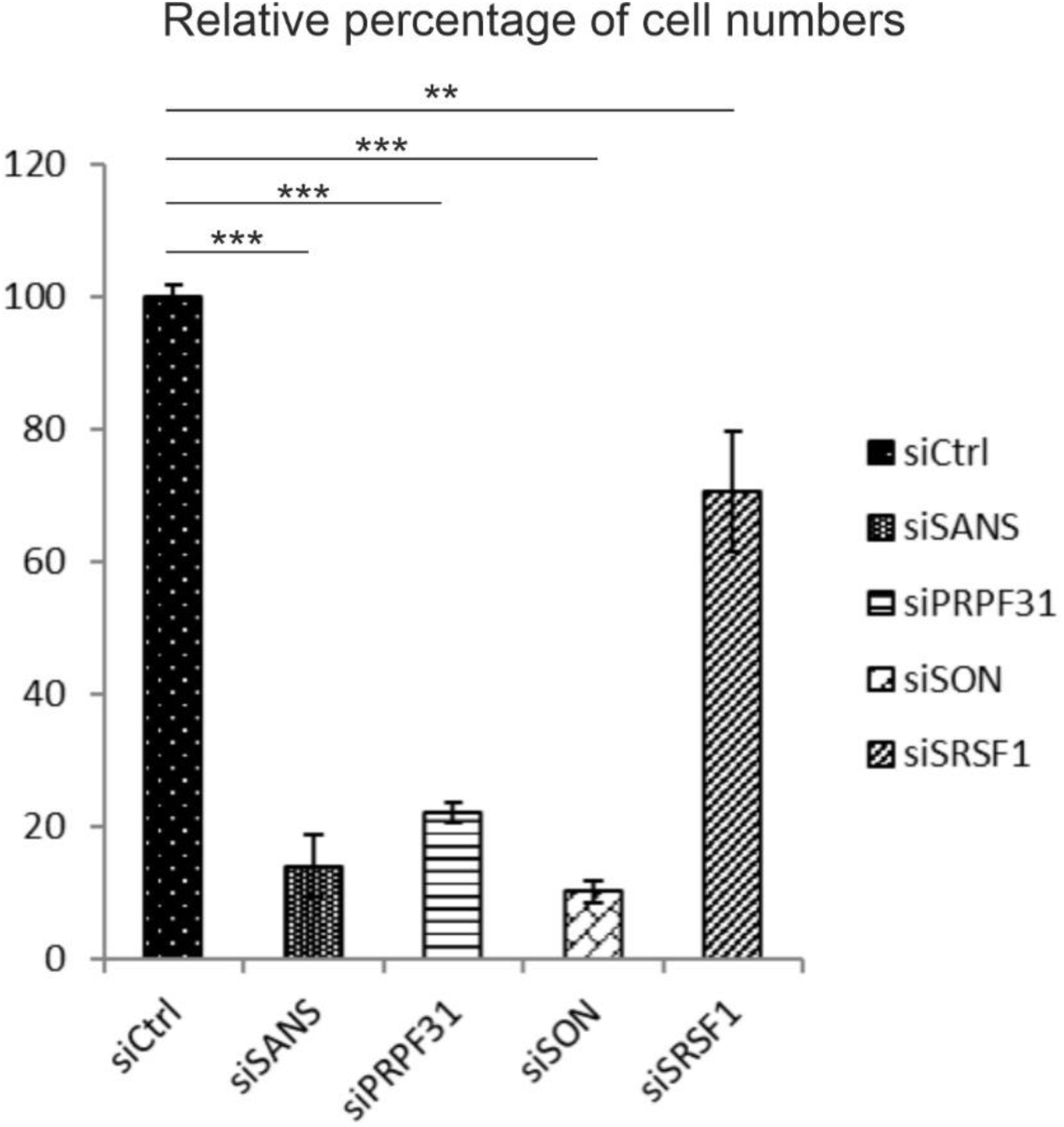
SANS depletion affects cell proliferation. WST-1 cell proliferation assay for cells depleted of SANS, SON, PRPF31 or SRSF1 by siRNA-mediated knockdown. siSANS, siSON and siPRPF31 reduce the proliferation rate by 80% when compared with control (siCtrl). In contrast, siSRSF1 reduces the proliferation rate only by less than 20%. ***: p<0.001, **: p<0.005.

## References

1. Papasaikas, P., Tejedor, J.R., Vigevani, L. and Valcarcel, J. (2015) Functional splicing network reveals extensive regulatory potential of the core spliceosomal machinery. Molecular cell, 57, 7–22.

2. Will, C.L. and Luhrmann, R. (2011) Spliceosome structure and function. Cold Spring Harbor perspectives in biology, 3.

3. Wahl, M.C., Will, C.L. and Luhrmann, R. (2009) The spliceosome: design principles of a dynamic RNP machine. Cell, 136, 701–718.

4. Novotny, I., Malinova, A., Stejskalova, E., Mateju, D., Klimesova, K., Roithova, A., Sveda, M., Knejzlik, Z. and Stanek, D. (2015) SART3-Dependent Accumulation of Incomplete Spliceosomal snRNPs in Cajal Bodies. Cell reports.

5. Michaud, S. and Reed, R. (1991) An ATP-independent complex commits pre-mRNA to the mammalian spliceosome assembly pathway. Genes & development, 5, 2534–2546.

6. Fischer, U., Englbrecht, C. and Chari, A. (2011) Biogenesis of spliceosomal small nuclear ribonucleoproteins. Wiley interdisciplinary reviews. RNA, 2, 718–731.

7. Roithova, A., Klimesova, K., Panek, J., Will, C.L., Luhrmann, R., Stanek, D. and Girard, C. (2018) The Sm-core mediates the retention of partially-assembled spliceosomal snRNPs in Cajal bodies until their full maturation. Nucleic acids research, 46, 3774–3790.

8. Stanek, D. (2017) Cajal bodies and snRNPs - friends with benefits. RNA biology, 14, 671–679.

9. Neugebauer, K.M. (2017) Special focus on the Cajal Body. RNA Biol, 14, 669–670.

10. Mao, Y.S., Zhang, B. and Spector, D.L. (2011) Biogenesis and function of nuclear bodies. Trends in genetics : TIG, 27, 295–306.

11. Galganski, L., Urbanek, M.O. and Krzyzosiak, W.J. (2017) Nuclear speckles: molecular organization, biological function and role in disease. Nucleic acids research, 45, 10350–10368.

12. Calarco, J.A., Saltzman, A.L., Ip, J.Y. and Blencowe, B.J. (2007) Technologies for the global discovery and analysis of alternative splicing. Advances in experimental medicine and biology, 623, 64–84.

13. Daguenet, E., Dujardin, G. and Valcarcel, J. (2015) The pathogenicity of splicing defects: mechanistic insights into pre-mRNA processing inform novel therapeutic approaches. EMBO reports, 16, 1640–1655.

14. Harbour, J.W. (2013) Genomic, prognostic, and cell-signaling advances in uveal melanoma. American Society of Clinical Oncology educational book. American Society of Clinical Oncology. Meeting, 388–391.

15. Quesada, V., Ramsay, A.J. and Lopez-Otin, C. (2012) Chronic lymphocytic leukemia with SF3B1 mutation. The New England journal of medicine, 366, 2530.

16. Tanackovic, G., Ransijn, A., Ayuso, C., Harper, S., Berson, E.L. and Rivolta, C. (2011) A missense mutation in PRPF6 causes impairment of pre-mRNA splicing and autosomal-dominant retinitis pigmentosa. Am J Hum Genet, 88, 643–649.

17. Overlack, N., Kilic, D., Bauss, K., Marker, T., Kremer, H., van Wijk, E. and Wolfrum, U. (2011) Direct interaction of the Usher syndrome 1G protein SANS and myomegalin in the retina. Biochim Biophys Acta, 1813, 1883–1892.

18. Mathur, P. and Yang, J. (2015) Usher syndrome: Hearing loss, retinal degeneration and associated abnormalities. Biochim Biophys Acta, 1852, 406–420.

19. Wolfrum, U. (2011) In Ahuja, S. (ed.), Usher Syndrome. Nova Science Publishers, Inc., pp. 51–73.

20. Liu, X., Vansant, G., Udovichenko, I.P., Wolfrum, U. and Williams, D.S. (1997) Myosin VIIa, the product of the Usher 1B syndrome gene, is concentrated in the connecting cilia of photoreceptor cells. Cell Motil Cytoskeleton, 37, 240–252.

21. Bauss, K., Knapp, B., Jores, P., Roepman, R., Kremer, H., Wijk, E.V., Marker, T. and Wolfrum, U. (2014) Phosphorylation of the Usher syndrome 1G protein SANS controls Magi2-mediated endocytosis. Human molecular genetics, 23, 3923–3942.

22. Jansen, F., Kalbe, B., Scholz, P., Mikosz, M., Wunderlich, K.A., Kurtenbach, S., Nagel-Wolfrum, K., Wolfrum, U., Hatt, H. and Osterloh, S. (2016) Impact of the Usher syndrome on olfaction. Human molecular genetics, 25, 524–533.

23. Bujakowska, K.M., Liu, Q. and Pierce, E.A. (2017) Photoreceptor Cilia and Retinal Ciliopathies. Cold Spring Harb Perspect Biol.

24. May-Simera, H., Nagel-Wolfrum, K. and Wolfrum, U. (2017) Cilia - The sensory antennae in the eye. Progress in retinal and eye research, 60, 144–180.

25. Chaki, M., Airik, R., Ghosh, A.K., Giles, R.H., Chen, R., Slaats, G.G., Wang, H., Hurd, T.W., Zhou, W., Cluckey, A. et al. (2012) Exome capture reveals ZNF423 and CEP164 mutations, linking renal ciliopathies to DNA damage response signaling. Cell, 150, 533–548.

26. Choi, H.J., Lin, J.R., Vannier, J.B., Slaats, G.G., Kile, A.C., Paulsen, R.D., Manning, D.K., Beier, D.R., Giles, R.H., Boulton, S.J. et al. (2013) NEK8 links the ATR-regulated replication stress response and S phase CDK activity to renal ciliopathies. Molecular cell, 51, 423–439.

27. Attanasio, M. (2015) Ciliopathies and DNA damage: an emerging nexus. Current opinion in nephrology and hypertension, 24, 366–370.

28. Gascue, C., Tan, P.L., Cardenas-Rodriguez, M., Libisch, G., Fernandez-Calero, T., Liu, Y.P., Astrada, S., Robello, C., Naya, H., Katsanis, N. et al. (2012) Direct role of Bardet-Biedl syndrome proteins in transcriptional regulation. Journal of cell science, 125, 362–375.

29. Sorusch, N., Bauss, K., Plutniok, J., Samanta, A., Knapp, B., Nagel-Wolfrum, K. and Wolfrum, U. (2017) Characterization of the ternary Usher syndrome SANS/ush2a/whirlin protein complex. Human molecular genetics, 26, 1157–1172.

30. Adato, A., Michel, V., Kikkawa, Y., Reiners, J., Alagramam, K.N., Weil, D., Yonekawa, H., Wolfrum, U., El-Amraoui, A. and Petit, C. (2005) Interactions in the network of Usher syndrome type 1 proteins. Human molecular genetics, 14, 347–356.

31. Lefevre, G., Michel, V., Weil, D., Lepelletier, L., Bizard, E., Wolfrum, U., Hardelin, J.P. and Petit, C. (2008) A core cochlear phenotype in USH1 mouse mutants implicates fibrous links of the hair bundle in its cohesion, orientation and differential growth. Development, 135, 1427–1437.

32. Caberlotto, E., Michel, V., Foucher, I., Bahloul, A., Goodyear, R.J., Pepermans, E., Michalski, N., Perfettini, I., Alegria-Prevot, O., Chardenoux, S. et al. (2011) Usher type 1G protein sans is a critical component of the tip-link complex, a structure controlling actin polymerization in stereocilia. Proc Natl Acad Sci U S A, 108, 5825–5830.

33. He, Y., Li, J. and Zhang, M. (2019) Myosin VII, USH1C, and ANKS4B or USH1G Together Form Condensed Molecular Assembly via Liquid-Liquid Phase Separation. Cell reports, 29, 974–986 e974.

34. Sahly, I., Dufour, E., Schietroma, C., Michel, V., Bahloul, A., Perfettini, I., Pepermans, E., Estivalet, A., Carette, D., Aghaie, A. et al. (2012) Localization of Usher 1 proteins to the photoreceptor calyceal processes, which are absent from mice. J Cell Biol, 199, 381–399.

35. Maerker, T., van Wijk, E., Overlack, N., Kersten, F.F., McGee, J., Goldmann, T., Sehn, E., Roepman, R., Walsh, E.J., Kremer, H. et al. (2008) A novel Usher protein network at the periciliary reloading point between molecular transport machineries in vertebrate photoreceptor cells. Human molecular genetics, 17, 71–86.

36. Sorusch, N., Yildirim, A., Knapp, B., Janson, J., Fleck, W., Scharf, C. and Wolfrum, U. (2019) SANS (USH1G) Molecularly Links the Human Usher Syndrome Protein Network to the Intraflagellar Transport Module by Direct Binding to IFT-B Proteins. Frontiers in cell and developmental biology, 7, 216.

37. Overlack, N., Maerker, T., Latz, M., Nagel-Wolfrum, K. and Wolfrum, U. (2008) SANS (USH1G) expression in developing and mature mammalian retina. Vision research, 48, 400–412.

38. Fabrizio, P., Laggerbauer, B., Lauber, J., Lane, W.S. and Luhrmann, R. (1997) An evolutionarily conserved U5 snRNP-specific protein is a GTP-binding factor closely related to the ribosomal translocase EF-2. The EMBO journal, 16, 4092–4106.

39. Fabrizio, P., Dannenberg, J., Dube, P., Kastner, B., Stark, H., Urlaub, H. and Luhrmann, R. (2009) The evolutionarily conserved core design of the catalytic activation step of the yeast spliceosome. Molecular cell, 36, 593–608.

40. Schindelin, J., Rueden, C.T., Hiner, M.C. and Eliceiri, K.W. (2015) The ImageJ ecosystem: An open platform for biomedical image analysis. Molecular reproduction and development, 82, 518–529.

41. Schindelin, J., Arganda-Carreras, I., Frise, E., Kaynig, V., Longair, M., Pietzsch, T., Preibisch, S., Rueden, C., Saalfeld, S., Schmid, B. et al. (2012) Fiji: an open-source platform for biological-image analysis. Nature methods, 9, 676–682.

42. Sedmak, T. and Wolfrum, U. (2010) Intraflagellar transport molecules in ciliary and nonciliary cells of the retina. J Cell Biol, 189, 171–186.

43. Sedmak, T., Sehn, E. and Wolfrum, U. (2009) Immunoelectron microscopy of vesicle transport to the primary cilium of photoreceptor cells. Methods Cell Biol, 94, 259–272.

44. Schaffert, N., Hossbach, M., Heintzmann, R., Achsel, T. and Luhrmann, R. (2004) RNAi knockdown of hPrp31 leads to an accumulation of U4/U6 di-snRNPs in Cajal bodies. The EMBO journal, 23, 3000–3009.

45. Taneja, K.L., Lifshitz, L.M., Fay, F.S. and Singer, R.H. (1992) Poly(A) RNA codistribution with microfilaments: evaluation by in situ hybridization and quantitative digital imaging microscopy. The Journal of cell biology, 119, 1245–1260.

46. Makarova, O.V., Makarov, E.M., Liu, S., Vornlocher, H.P. and Luhrmann, R. (2002) Protein 61K, encoded by a gene (PRPF31) linked to autosomal dominant retinitis pigmentosa, is required for U4/U6*U5 tri-snRNP formation and pre-mRNA splicing. The EMBO journal, 21, 1148–1157.

47. Agafonov, D.E., Kastner, B., Dybkov, O., Hofele, R.V., Liu, W.T., Urlaub, H., Luhrmann, R. and Stark, H. (2016) Molecular architecture of the human U4/U6.U5 tri-snRNP. Science, 351, 1416–1420.

48. Buskin, A., Zhu, L., Chichagova, V., Basu, B., Mozaffari-Jovin, S., Dolan, D., Droop, A., Collin, J., Bronstein, R., Mehrotra, S. et al. (2018) Disrupted alternative splicing for genes implicated in splicing and ciliogenesis causes PRPF31 retinitis pigmentosa. Nature communications, 9, 4234.

49. Golas, M.M., Sander, B., Will, C.L., Luhrmann, R. and Stark, H. (2003) Molecular architecture of the multiprotein splicing factor SF3b. Science, 300, 980–984.

50. Cretu, C., Schmitzova, J., Ponce-Salvatierra, A., Dybkov, O., De Laurentiis, E.I., Sharma, K., Will, C.L., Urlaub, H., Luhrmann, R. and Pena, V. (2016) Molecular Architecture of SF3b and Structural Consequences of Its Cancer-Related Mutations. Molecular cell, 64, 307–319.

51. Prilusky, J., Felder, C.E., Zeev-Ben-Mordehai, T., Rydberg, E.H., Man, O., Beckmann, J.S., Silman, I. and Sussman, J.L. (2005) FoldIndex: a simple tool to predict whether a given protein sequence is intrinsically unfolded. Bioinformatics, 21, 3435–3438.

52. Linding, R., Jensen, L.J., Diella, F., Bork, P., Gibson, T.J. and Russell, R.B. (2003) Protein disorder prediction: implications for structural proteomics. Structure, 11, 1453–1459.

53. Wright, P.E. and Dyson, H.J. (2015) Intrinsically disordered proteins in cellular signalling and regulation. Nature reviews. Molecular cell biology, 16, 18–29.

54. Nguyen Ba, A.N., Pogoutse, A., Provart, N. and Moses, A.M. (2009) NLStradamus: a simple Hidden Markov Model for nuclear localization signal prediction. BMC bioinformatics, 10, 202.

55. Brameier, M., Krings, A. and MacCallum, R.M. (2007) NucPred--predicting nuclear localization of proteins. Bioinformatics, 23, 1159–1160.

56. Fei, J., Jadaliha, M., Harmon, T.S., Li, I.T.S., Hua, B., Hao, Q., Holehouse, A.S., Reyer, M., Sun, Q., Freier, S.M. et al. (2017) Quantitative analysis of multilayer organization of proteins and RNA in nuclear speckles at super resolution. Journal of cell science, 130, 4180–4192.

57. Cremer, T. and Cremer, C. (2001) Chromosome territories, nuclear architecture and gene regulation in mammalian cells. Nature reviews. Genetics, 2, 292–301.

58. Spector, D.L. and Lamond, A.I. (2011) Nuclear speckles. Cold Spring Harbor perspectives in biology, 3.

59. Schutze, T., Ulrich, A.K., Apelt, L., Will, C.L., Bartlick, N., Seeger, M., Weber, G., Luhrmann, R., Stelzl, U. and Wahl, M.C. (2016) Multiple protein-protein interactions converging on the Prp38 protein during activation of the human spliceosome. Rna, 22, 265–277.

60. Liu, S., Rauhut, R., Vornlocher, H.P. and Luhrmann, R. (2006) The network of protein-protein interactions within the human U4/U6.U5 tri-snRNP. Rna, 12, 1418–1430.

61. Huang da, W., Sherman, B.T. and Lempicki, R.A. (2009) Systematic and integrative analysis of large gene lists using DAVID bioinformatics resources. Nature protocols, 4, 44–57.

62. Sharma, A., Takata, H., Shibahara, K., Bubulya, A. and Bubulya, P.A. (2010) Son is essential for nuclear speckle organization and cell cycle progression. Molecular biology of the cell, 21, 650–663.

63. Lu, X., Goke, J., Sachs, F., Jacques, P.E., Liang, H., Feng, B., Bourque, G., Bubulya, P.A. and Ng, H.H. (2013) SON connects the splicing-regulatory network with pluripotency in human embryonic stem cells. Nature cell biology, 15, 1141–1152.

64. Reiners, J., Reidel, B., El-Amraoui, A., Boeda, B., Huber, I., Petit, C. and Wolfrum, U. (2003) Differential distribution of harmonin isoforms and their possible role in Usher-1 protein complexes in mammalian photoreceptor cells. Invest Ophthalmol Vis Sci, 44, 5006–5015.

65. Chen, Z.Y., Hasson, T., Kelley, P.M., Schwender, B.J., Schwartz, M.F., Ramakrishnan, M., Kimberling, W.J., Mooseker, M.S. and Corey, D.P. (1996) Molecular cloning and domain structure of human myosin-VIIa, the gene product defective in Usher syndrome 1B. Genomics, 36, 440–448.

66. Papal, S., Cortese, M., Legendre, K., Sorusch, N., Dragavon, J., Sahly, I., Shorte, S., Wolfrum, U., Petit, C. and El-Amraoui, A. (2013) The giant spectrin betaV couples the molecular motors to phototransduction and Usher syndrome type I proteins along their trafficking route. Human molecular genetics, 22, 3773–3788.

67. Sorusch, N., Wunderlich, K., Bauss, K., Nagel-Wolfrum, K. and Wolfrum, U. (2014) Usher syndrome protein network functions in the retina and their relation to other retinal ciliopathies. Advances in experimental medicine and biology, 801, 527–533.

68. van Wijk, E., Kersten, F.F., Kartono, A., Mans, D.A., Brandwijk, K., Letteboer, S.J., Peters, T.A., Marker, T., Yan, X., Cremers, C.W. et al. (2009) Usher syndrome and Leber congenital amaurosis are molecularly linked via a novel isoform of the centrosomal ninein-like protein. Human molecular genetics, 18, 51–64.

69. De Biasio, A., Ibanez de Opakua, A., Cordeiro, T.N., Villate, M., Merino, N., Sibille, N., Lelli, M., Diercks, T., Bernado, P. and Blanco, F.J. (2014) p15PAF is an intrinsically disordered protein with nonrandom structural preferences at sites of interaction with other proteins. Biophysical journal, 106, 865–874.

70. Protter, D.S.W., Rao, B.S., Van Treeck, B., Lin, Y., Mizoue, L., Rosen, M.K. and Parker, R. (2018) Intrinsically Disordered Regions Can Contribute Promiscuous Interactions to RNP Granule Assembly. Cell reports, 22, 1401–1412.

71. Alberti, S. and Dormann, D. (2019) Liquid-Liquid Phase Separation in Disease. Annual review of genetics, 53, 171–194.

72. Banani, S.F., Lee, H.O., Hyman, A.A. and Rosen, M.K. (2017) Biomolecular condensates: organizers of cellular biochemistry. Nature reviews. Molecular cell biology, 18, 285–298.

73. Banani, S.F., Rice, A.M., Peeples, W.B., Lin, Y., Jain, S., Parker, R. and Rosen, M.K. (2016) Compositional Control of Phase-Separated Cellular Bodies. Cell, 166, 651–663.

74. Girard, C., Will, C.L., Peng, J., Makarov, E.M., Kastner, B., Lemm, I., Urlaub, H., Hartmuth, K. and Luhrmann, R. (2012) Post-transcriptional spliceosomes are retained in nuclear speckles until splicing completion. Nature communications, 3, 994.

75. Novotny, I., Blazikova, M., Stanek, D., Herman, P. and Malinsky, J. (2011) In vivo kinetics of U4/U6.U5 tri-snRNP formation in Cajal bodies. Molecular biology of the cell, 22, 513–523.

76. Ahn, E.Y., DeKelver, R.C., Lo, M.C., Nguyen, T.A., Matsuura, S., Boyapati, A., Pandit, S., Fu, X.D. and Zhang, D.E. (2011) SON controls cell-cycle progression by coordinated regulation of RNA splicing. Molecular cell, 42, 185–198.

77. Boesler, C., Rigo, N., Anokhina, M.M., Tauchert, M.J., Agafonov, D.E., Kastner, B., Urlaub, H., Ficner, R., Will, C.L. and Luhrmann, R. (2016) A spliceosome intermediate with loosely associated tri-snRNP accumulates in the absence of Prp28 ATPase activity. Nature communications, 7, 11997.

78. Zou, J., Chen, Q., Almishaal, A., Mathur, P.D., Zheng, T., Tian, C., Zheng, Q.Y. and Yang, J. (2017) The roles of USH1 proteins and PDZ domain-containing USH proteins in USH2 complex integrity in cochlear hair cells. Human molecular genetics, 26, 624–636.

79. Matera, A.G. and Wang, Z. (2014) A day in the life of the spliceosome. Nature reviews. Molecular cell biology, 15, 108–121.

80. Wheway, G., Schmidts, M., Mans, D.A., Szymanska, K., Nguyen, T.M., Racher, H., Phelps, I.G., Toedt, G., Kennedy, J., Wunderlich, K.A. et al. (2015) An siRNA-based functional genomics screen for the identification of regulators of ciliogenesis and ciliopathy genes. Nature cell biology, 17, 1074–1087.

81. Williams, D.S. (2008) Usher syndrome: animal models, retinal function of Usher proteins, and prospects for gene therapy. Vision research, 48, 433–441.

82. Graziotto, J.J., Farkas, M.H., Bujakowska, K., Deramaudt, B.M., Zhang, Q., Nandrot, E.F., Inglehearn, C.F., Bhattacharya, S.S. and Pierce, E.A. (2011) Three gene-targeted mouse models of RNA splicing factor RP show late-onset RPE and retinal degeneration. Investigative ophthalmology & visual science, 52, 190–198.

83. Farkas, M.H., Grant, G.R., White, J.A., Sousa, M.E., Consugar, M.B. and Pierce, E.A. (2013) Transcriptome analyses of the human retina identify unprecedented transcript diversity and 3.5 Mb of novel transcribed sequence via significant alternative splicing and novel genes. BMC Genomics, 14, 486.

84. Zambelli, F., Pavesi, G., Gissi, C., Horner, D.S. and Pesole, G. (2010) Assessment of orthologous splicing isoforms in human and mouse orthologous genes. BMC genomics, 11, 534.

85. Xu, M., Xie, Y.A., Abouzeid, H., Gordon, C.T., Fiorentino, A., Sun, Z., Lehman, A., Osman, I.S., Dharmat, R., Riveiro-Alvarez, R. et al. (2017) Mutations in the Spliceosome Component CWC27 Cause Retinal Degeneration with or without Additional Developmental Anomalies. American journal of human genetics, 100, 592–604.

